# Systemic impact of the expression of the mitochondrial alternative oxidase on *Drosophila* development

**DOI:** 10.1101/2021.09.24.461559

**Authors:** André F. Camargo, Sina Saari, Geovana S. Garcia, Marina M. Chioda, Murilo F. Othonicar, Ailton A. Martins, Gabriel Hayashi, Johanna ten Hoeve, Howard T. Jacobs, Daniel G. Pinheiro, Eric Dufour, Marcos T. Oliveira

## Abstract

Despite the beneficial effects of xenotopically expressing the mitochondrial alternative oxidase AOX from *Ciona intestinalis* in mammalian and insect models, important detrimental outcomes have also been reported, raising concerns regarding its potential deployment as a therapeutic enzyme for human mitochondrial diseases. Because of its non-protonmotive terminal oxidase activity, AOX can bypass the cytochrome segment of the respiratory chain whilst not contributing to mitochondrial ATP synthesis. We have previously shown that pupal lethality occurs when AOX-expressing *Drosophila* larvae are cultured on a low-nutrient diet, indicating that AOX can perturb normal metabolism during development. Here, combined omics analyses revealed multiple correlates of this diet-dependent lethality, including a general alteration of larval amino acid and lipid metabolism, functional and morphological changes to the larval digestive tract, and a drastic decrease in larval biomass accumulation. Pupae at the pre-lethality stage presented a general downregulation of mitochondrial metabolism and a signature of starvation and deregulated signaling. AOX-induced lethality was partially rescued when the low-nutrient diet was supplemented with tryptophan and/or methionine, but not with proline and/or glutamate, strongly suggesting perturbation of one-carbon metabolism. The developmental dependence on tryptophan and/or methionine, associated with elevated levels of lactate dehydrogenase, 2-hydroxyglutarate, choline-containing metabolites and breakdown products of membrane phospholipids, indicates that AOX expression promotes tissue proliferation and larval growth, but this is ultimately limited by energy dissipation due to partial mitochondrial uncoupling. We speculate that the combination of dietary interventions and AOX expression might, nevertheless, be useful for the metabolic regulation of proliferative tissues, such as tumors.

## Introduction

Although mitochondria have many important roles in eukaryotic cells, the bulk production of ATP via oxidative phosphorylation (OXPHOS) is traditionally viewed as one of their main contributing processes for cellular function. Several mitochondrial inner membrane-embedded enzymes, such as NADH:ubiquinone oxidoreductase (complex I, CI), succinate dehydrogenase (complex II, CII), and mitochondrial glycerol-3-phosphate dehydrogenase (mGPDH), among others, may convergently initiate OXPHOS by catalyzing the reduction of coenzyme Q (CoQ/CoQH_2_). Because they re-oxidize key metabolites and electron carriers located in the mitochondrial matrix and outside mitochondria, these dehydrogenases may directly control the metabolism of nutrients such as monosaccharides, amino acids, and fatty acids. Starting a serial chain of redox reactions that we herein refer to as the cytochrome segment of the respiratory chain, the cytochrome *bc_1_* complex (complex III, CIII) oxidizes CoQH_2_, reducing cytochrome *c*, which in turn is re-oxidized by the cytochrome *c* oxidase (complex IV, CIV) in a reaction that uses molecular oxygen (O_2_) as the final electron acceptor, releasing H_2_O. This electron transfer system (ETS) formed by the aforementioned dehydrogenases combined with the cytochrome segment is paired with the pumping of protons from the mitochondrial matrix into the intermembrane space. The energy of the electrochemical gradient thus formed is eventually used by the ATP synthase complex to phosphorylate ADP, which returns protons to the mitochondrial matrix.

The electrochemical gradient, the production of ATP, and the reoxidation of the intermediate electron carriers are crucial for the regulation of cellular metabolism. OXPHOS function directly modulates the catabolic and anabolic functions of the tricarboxylic acid (TCA) cycle, and may have various consequences depending on factors such as tissue/cell type, stage of development, temperature and/or nutritional availability. For example, in proliferative tissues such as in tumors or in the growing *Drosophila* larva, a significant fraction of the TCA cycle intermediates serves as precursors for the synthesis of lipids, nucleotides, carbohydrates and proteins, in a process referred to as cataplerosis (1). Although cataplerosis appears to be the most important function of mitochondria during biomass accumulation, it is strongly impaired if the reoxidation of NADH and/or CoQH_2_ by the ETS is compromized (2, 3).

Protists, fungi, plants, and most metazoans, but not vertebrates or insects, are endowed with a branched OXPHOS system involving a second terminal oxidase, the non-protonmotive alternative oxidase (AOX) (4). AOX activity bypasses CIII/CIV, directly coupling O_2_ reduction with CoQH_2_ oxidation. Although AOX activity may decrease mitochondrial ATP synthesis, it maintains electron flow and cellular redox potential, and decreases reactive oxygen species (ROS) production caused by OXPHOS dysfunction (e.g. stress, overload or blockage) (3,5–7). In humans, severe OXPHOS dysfunction is related to a variety of mitochondrial diseases, whose onset and severity vary broadly, with very limited effective treatments (8). The premise that AOX activity would be beneficial for higher animals with OXPHOS dysfunction has been successfully tested in several models (9–14). In addition to supporting respiration resistant to inhibitors of CIII and CIV, AOX enabled complete or partial recovery of deleterious phenotypes caused by mutations in subunits of the OXPHOS complexes (3,10,15–17) and is under consideration as a therapy enzyme (18, 19).

Nevertheless, AOX expression studies in model organisms have been limited in regard to the alterations it may cause to normal metabolism and physiology, especially considering that AOX activity could partially uncouple mitochondria and generate excess heat. We have been using *Drosophila melanogaster* lines expressing AOX from the tunicate *Ciona intestinalis* (Ascidiacea) to assess possible limitations in the use of this enzyme in higher animals. We have shown that AOX expression causes a dose-dependent depletion of mature spermatozoids, which leads to a dramatic disadvantage for AOX-expressing males in sperm competition assays (20). We also recently reported that AOX expression heavily compromises adult eclosion when the flies are cultured on a low nutrient (LN), but not on standard laboratory (SD) diet (21). Our SD diet contains 1.5% sucrose (w/v), 3% glucose, 1.5% maize flour, 1% wheat germ, 1% soy flour, 3% molasses, and 3.5% yeast extract, whereas in the LN diet, the only source of nutrients is 3.5% yeast extract (w/v), which is normally sufficient for the *Drosophila* larvae to reach the adult stage. Supplementation of the LN diet showed that nutritionally and structurally complex ingredients of the SD diet, such as maize flour, wheat germ or molasses, could rescue the lethality of AOX-expressing flies, but not simple compounds like sugars, vitamins and minerals, nor additional yeast extract (22). Mass spectrometry revealed a complex array of metabolites in a fraction of molasses that is rich in intermediates of the TCA cycle, among other molecules, but no compound that has been tested to date could individually rescue the lethality of AOX-expressing flies cultured on LN. This suggests that AOX may cause complex metabolic changes, which could explain why the transgene is unable to rescue deleterious phenotypes of some well-established mitochondrial mutants (23–26), and yet it was effective against genetic defects with no apparent direct link to mitochondria (27).

Here we applied a multi-omics approach to gain more insight into the molecular basis of the lethal developmental interaction between AOX expression and LN diet. Our data suggests that AOX expression and LN diet have opposite effects on larval amino acid metabolism. AOX expression is also accompanied by increased larval feeding behavior, functional and morphological alterations of the larval intestines, impaired nutrient storage in the larvae, and signs of starvation and deregulated signaling in the pupae. Pupal lethality was partially rescued by LN supplementation of the feeding larvae with methionine or tryptophan, suggesting an important role for one-carbon metabolism in the adaptive processes that allow AOX-expressing flies to complete development. We discuss the implications of our findings for tissue growth and AOX therapy.

## Results and Discussion

### AOX expression causes systemic developmental failure on low nutrient diet

In a previously published study (21), we showed that the LN diet-dependent pupal lethality was achieved by high levels of ubiquitously-driven AOX expression. Here, we analyzed additional *UAS-AOX* lines that, when crossed with the *daGAL4* driver line, provided flies with varying levels of ubiquitous AOX expression, assigned as “weak”, “intermediate” and “strong”. We confirmed that in LN diet pupal lethality only occurs with strong expression of AOX (Figure 1A), suggesting that the phenotype is dependent on a threshold effect. We also verified that the LN diet does not alter AOX protein or transcript levels (Figure 1B, C). From this point on, unless otherwise stated, we continued to analyze only the flies with strong AOX expression (*daGAL4*-driven progeny of the line *UAS-AOX_F6_* (10)), referring to them simply as AOX-expressing flies.

**Figure 1.**
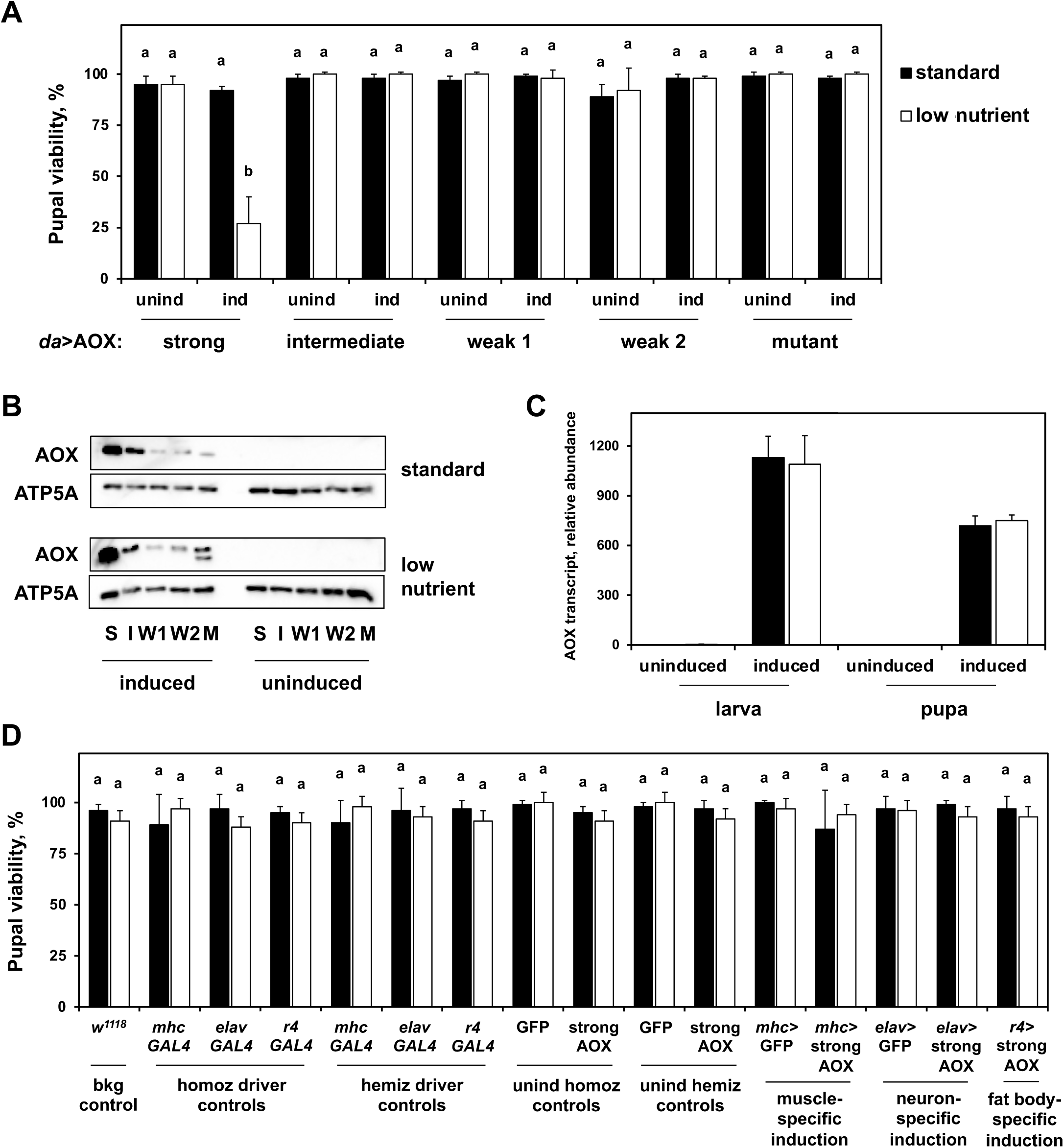
Pupal lethality is achieved via interaction between low nutrient diet and strong ubiquitous expression of AOX. A, Pupal viability (adult eclosion percentage) of flies with ubiquitous induction (ind) or not (unind) of AOX expression at the indicated levels. AOX expression was accomplished by crossing the homozygous *daGAL4* line with homozygous *UAS-AOX_F6_* (10) for strong levels, *UAS-AOX_8.1_; UAS-AOX_7.1_* for intermediate levels, *UAS-AOX_8.1_* and *UAS-AOX_7.1_* (16) for weak 1 and 2 levels, and *UAS-mutAOX_2nd_; UAS-mutAOX_3rd_* for the catalytically-inactive mutant form of AOX (16). Uninduced controls were progeny of the same *UAS* lines and *w_1118_*. B, Immunoblot showing levels in whole body samples of a mitochondrial protein marker, ATP5A, and AOX when its expression was ubiquitously induced at strong (S), intermediate (I) or weak (W1, W2) levels on the indicated diet, using the same lines as in A. M, mutant AOX line, which showed expression lower than expected and partial degradation of the mutant protein. C, AOX transcript levels extracted from the RNA-seq data of whole-body samples of flies with strong ubiquitous expression of AOX (progeny of *UAS-AOX_F6_* and *daGAL4*) and uninduced controls (progeny of *UAS-AOX_F6_* and *w_1118_*) at the indicated developmental stages, cultured on the indicated diet. Data normalized by the read counts in uninduced controls cultured on standard diet, arbitrarily set to 1.0. D, Pupal viability of control flies, and flies with tissue-specific induction or not (unind hemiz controls) of strong expression of AOX in muscles, neurons or fat body, when cultured at the indicated diet. bkg, genetic background; homoz, homozygous; hemiz, hemizygous; unind, uninduced. The homoz driver and unind homoz controls represent, respectively, the *GAL4* driver and *UAS* lines. The hemiz driver and unind hemiz controls represent, respectively, flies with a single copy of the transgenic *GAL4* and *UAS* constructs, i.e., progeny of the cross between the homozygous lines with *w_1118_*. Tissue-specific induction was accomplished by crossing homozygous *GAL4* with *UAS* lines, resulting in flies with a single hemizygous copy of each construct. GFP, *UAS-StingerGFP*; strong AOX, *UAS-AOX_F6_* (10). The data in A, C and D represent average of two or three experiments; error bars represent standard deviations. a and b represent significantly different statistical classes (one-way ANOVA, followed by the Tukey *post-hoc* test, p < 0.05).

We also drove expression of AOX in three larval tissues of major importance for whole-animal energy metabolism, to test if the lethality of the AOX-expressing flies in LN diet was due to tissue-specific effects. AOX expression in the larval musculature, the nervous system, or in the fat body, in combination with LN diet, had no effect on pupal viability (Figure 1D). The lethal phenotype is therefore the consequence of AOX expression in a less prominent tissue or a systemic effect initiated in multiple tissues.

### AOX expression induces functional and morphological changes in the larval digestive system

Given the impracticality of testing the many hundreds of tissue-restricted *GAL4* driver lines available, in all possible combinations, we instead performed deep RNA sequencing (RNA-seq) of whole individual specimens, with the aim of identifying general and tissue-specific transcriptional changes linked to the interaction between LN diet and AOX expression. Although this approach may not reveal spatial gene misexpression, we reasoned that it should enable us to develop a mechanistic hypothesis. We selected wandering third-instar (L3) larvae and pre-pharate pupae, the immediate post-feeding and pre-mortem stages, respectively. General analyses of the data either separately (Supplemental Tables S1-S6) or combined (Supplemental Tables S7-S8) showed that the transcriptomic changes from larva to pre-pharate pupa were as expected and previously described (28, 29). The pupae showed a general upregulation of mitochondrial and nervous-system genes, consistent with a developmental switch from a glycolytic to an OXPHOS-based metabolism (30, 31), and with the morphogenesis of the adult brain and eyes. We also observed downregulation of genes involved in proteolytic/autophagic processes, information pathways/cell cycle, sugar transport and metabolism, and genes encoding alkaline phosphatases, likely characterizing the end of the metamorphic phase. Remodeling of the epidermis/cuticle/exoskeleton is also apparent by the up and downregulation of different subsets of genes (Supplemental Tables S1-S8), most probably reflecting the cell migration events that ensure dorsal thoracic and abdominal midline closure (32).

Multifactorial analysis of data from larvae showed that the LN diet causes the largest change in the fly transcriptome, altering the levels of ∼10% of the 13809 unique transcripts identified (Supplemental Table S9). Enrichment analysis and functional annotation revealed that metabolic processes and nutrient transport were extensively affected by the LN diet, as expected, in addition to genes associated with sexual maturation, RNA binding, and protein folding (Supplemental Figure S1). AOX expression altered the levels of ∼2% of the identified transcripts (Supplemental Table S10), and the interaction between LN and AOX (LN-AOX) altered another ∼2% (Supplemental Table S11). In both cases, only a few clusters were assigned (Figure 2, Supplemental Figures S2-S3), the signaling and membrane protein clusters being the largest ones (see next sections for details).

**Figure 2.**
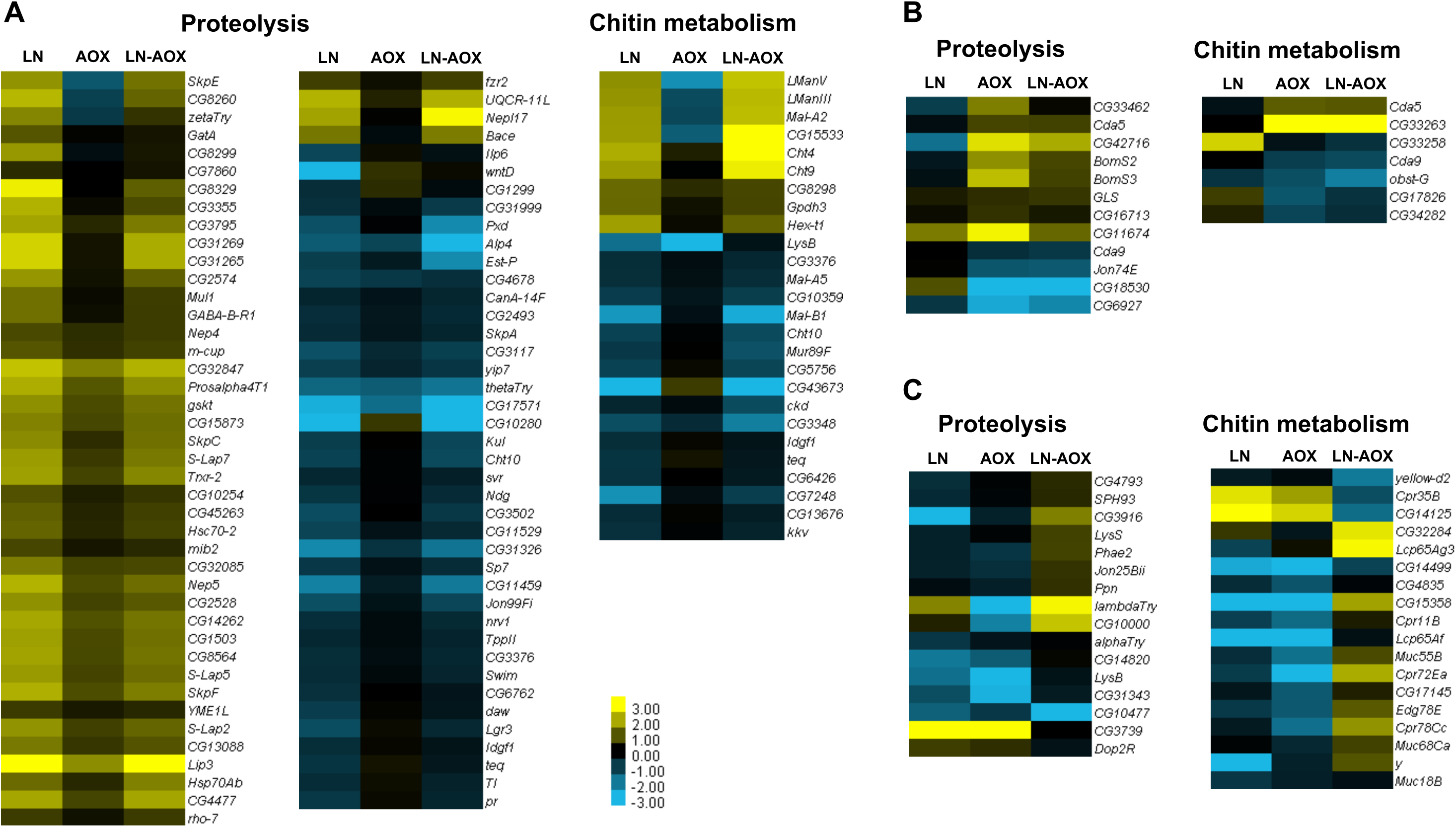
Transcripts involved with proteolysis and chitin metabolism are commonly affected by low nutrient (LN) diet and/or AOX expression in larvae. Heatmaps show the transcripts altered by LN diet (A), strong ubiquitous AOX expression (B), and the LN-AOX interaction (C). The columns LN, AOX and LN-AOX show log_2_(fold change) values of the indicated transcript in each condition in pairwise comparisons with control larvae cultured on SD diet. See Supplemental Figures S1-S3 for other transcript clusters, and Material and Methods for experimental details. Control larvae, progeny of *UAS-AOX^F6^* and *w^1118^*; AOX-expressing larvae, progeny of *UAS-AOX^F6^* and *daGAL4*.

We found two common gene clusters with a mixture of up and downregulated transcripts for LN diet, AOX expression, and LN-AOX interaction: proteolysis and chitin metabolism (Figure 2). Data from modENCODE (28, 33) deposited in Flybase (34) highlighted the tissues in which the transcripts from these two clusters are abundant. The proteolysis and chitin metabolism transcripts altered by the LN diet (Figure 2A) appear to be scattered among different organs (Supplemental Table S12), whereas the transcripts altered by AOX expression (Figure 2B) and LN-AOX interaction (Figure 2C) were most commonly found in the digestive system and salivary glands (Supplemental Tables S13 and S14). Interestingly, when analyzing the transcripts in all clusters (not only in the proteolysis and chitin metabolism clusters) in the AOX and the LN-AOX groups, we found that 63 and 58% are usually associated with digestive tissues, respectively. In LN diet, transcripts related to larval/adult digestion only account for ∼20% of total clustered transcripts (Supplemental Tables S12-S14).

The majority of genes in the proteolysis clusters we identified were endo/exo peptidases, one of the largest families of digestive enzymes in flies (35). Flies also express a variety of chitinases and glucanases that play a role in yeast cell digestion, which are especially important in the context of our LN “yeast extract only” diet (21). Chitin metabolism genes are also important for the development and function of the internal lining of the intestines, which consists primarily of epithelial cells containing chitin-based structures that are frequently remodeled to allow proper digestion and insect growth (36). Although the gastrointestinal tract of *Drosophila* is a complex organ with multiple cell types of different developmental origin (37), our data suggest that AOX expression alters larval digestion or nutrient absorption, before the animals enter the pupal stage. To test this hypothesis, we checked the intestinal morphology of the larvae, revealing that LN diet causes shortening of the intestines in the control flies, which was prevented by AOX expression (Figure 3A). We next tested whether tissue-restricted expression of AOX could reproduce the pupal lethality observed with the ubiquitous expression system, by the use of a GAL4 driver supporting expression principally in cells of the gut, as well as some other epithelia (38–40). However, pupal viability remained as high as in the controls (Figure 3B), indicating that the lethal phenotype is a consequence of AOX expression in one or more other tissues, and could be a systemic effect in which the digestive tract is significantly affected, perhaps as part of a compensatory physiological response.

**Figure 3.**
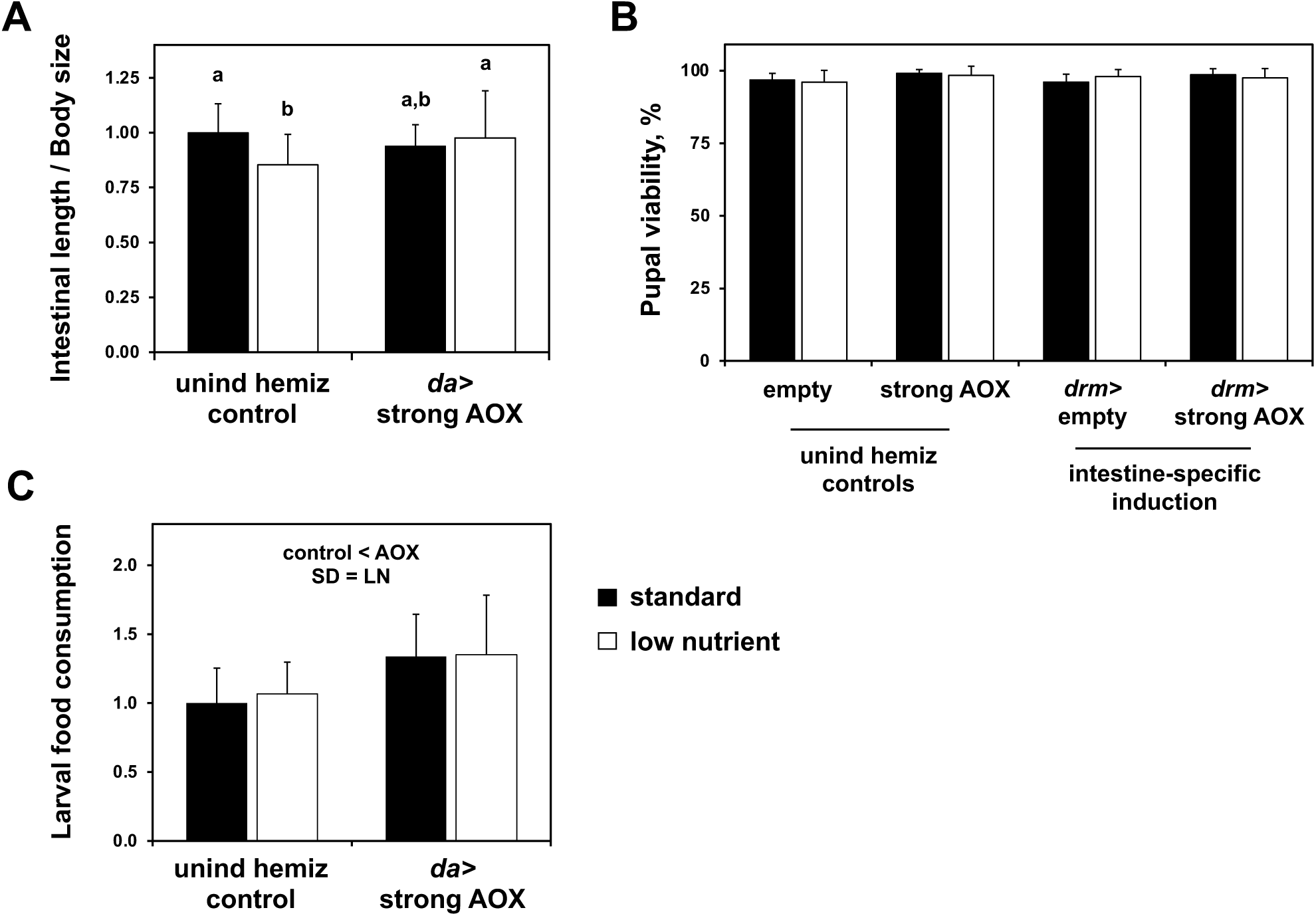
Impact of diet and AOX expression on larval intestines and food consumption. A, Intestinal length of larvae of the indicated genotypes cultured under the diets shown, normalized against intestinal length of uninduced controls cultured in SD diet. a and b represent significantly different classes (two-way ANOVA, followed by Tukey HSD *post-hoc* test, p < 0.05). B, Pupal viability of flies with or without AOX induction on the indicated diet, using *drmGAL4* to drive expression predominantly in the gut. The data represent averages of at least two independent experiments, and the error bars represent standard deviations. C, Food consumption normalized to value for uninduced control larvae cultured on standard diet. “control < AOX” indicates significant differences between control and AOX-expressing flies (two-way ANOVA, p < 0.05).

### AOX expression leads to decreased larval biomass accumulation

A compensatory response that changes the larval gut both functionally and morphologically suggests that AOX expression might disturb food consumption, absorption, and/or larval biomass accumulation. We first examined food consumption in the early L3 stage and observed that AOX-expressing larvae eat more, independently of diet (Figure 3C). However, the increased feeding behavior was not associated with gain in larval weight. Instead, although developing normally, AOX larvae cultured on SD diet had a body mass equivalent to that of control larvae on LN diet (i. e., decreased by ∼15%). The effects of the two conditions were additive, leading in the LN-AOX larvae to a ∼40% decreased body mass (Figure 4A-C), accompanied by a decrease in larval and pupal size (Supplemental Figure S4). Interestingly, we previously reported that adult flies expressing AOX with the same *UAS*/*GAL4* system, cultured on SD diet, present a more pronounced loss of weight as the flies age (10).

**Figure 4.**
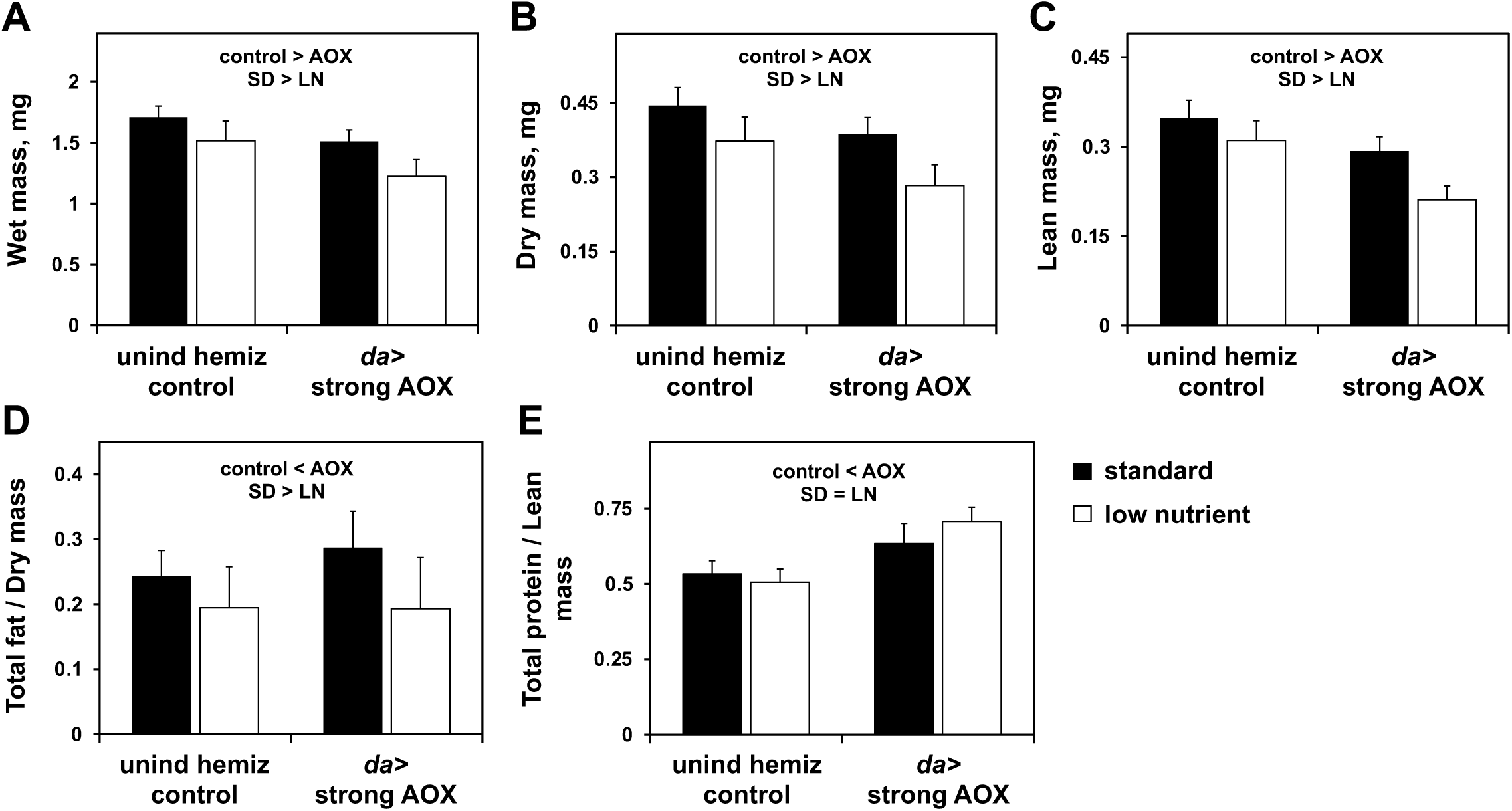
The combination of strong ubiquitous expression of AOX and low nutrient diet promotes a dramatic decrease in larval body mass. Wet (A), dry (B) and lean (C) masses and total (D) fat and (E) protein of L3 larvae of the indicated genotypes, cultured on diets as shown. unind hemiz control, progeny of *UAS-AOX^F6^* and *w^1118^*; *da*>strong AOX, progeny of *UAS-AOX^F6^* and *daGAL4*. “control > AOX” and “control < AOX” indicate significant differences between control and AOX-expressing flies; “SD > LN”, between diets (two-way ANOVA, p < 0.05).

Decreased larval biomass despite increased food uptake suggests metabolic wasting in AOX-expressing flies, particularly when cultured on LN diet. We previously observed that larval triglycerides were significantly lower in larvae cultured on LN, but AOX expression had no effect on the levels of these storage molecules (22). Because in *Drosophila* triglyceride levels increase as the larvae grow, and continue increasing during metamorphosis at the expense of membrane lipids (41), we quantitated total fat content in our larvae by ether extraction to derive an estimate for combined triglycerides, membrane lipids, and other lipids. The LN diet lowered total larval fat, which is consistent with the drop in triglycerides previously seen (22). In contrast, AOX-expressing larvae showed increased fat content, which is apparent only in SD diet (Figure 4D). Although how these alterations relate to fly survival remain to be addressed, it is noticeable that the LN diet causes a substantial change in lipid metabolism transcripts (Supplemental Figure S1), of which several fatty acid β-oxidation genes are downregulated (Supplemental Figure S5). In addition, both LN diet and LN-AOX caused significant changes in the levels of larval membrane proteins (Supplemental Figures S1 and S3), which are usually affected by lipid composition, and LN-AOX pupae showed elevated transcript levels for three enzymes involved in triglyceride biosynthesis (Supplemental Figure S5), consistent with a response to lipid deficiency during metamorphosis.

We also estimated the total content of larval protein, as this is another class of stored nutrient important for pupal development (42). AOX expression was associated with a 25-40% increase in total protein levels, irrespective of diet (Figure 4E). This is consistent with the increased feeding behavior (Figure 3C), as well as with the changes in proteolysis detected in the transcriptome (Figure 2) and in amino acid levels (see below). Taken together, our data indicates that AOX-expressing larvae accumulate more fat and protein for a given amount of larval mass, when cultured on SD diet. This may indicate a successful compensatory adaptation to decreased mitochondrial energy efficiency, achieved through increased nutrient absorption and storage in the larvae, allowing the flies to traverse metamorphosis and reach the adult stage, despite the energy dissipation caused by AOX activity. The LN diet, in which yeast extract is the sole nutrient source, appears to prevent this increase in fat but not in protein accumulation. We, therefore, speculate that the AOX-associated changes observed in the larval gut and in feeding behavior are likely due to altered nutrient absorption, geared to enabling larvae to accumulate sufficient nutrients for the completion of development. However, on LN diet, this is insufficient to compensate for the effects of AOX, leading to metabolic imbalance in larvae and wasting in pupae.

### The interaction between AOX expression and low nutrient diet leads to developmental arrest due to pupal starvation

We next investigated the pre-pharate transcriptional landscape to gain insight into the molecular events immediately preceding the death of LN-AOX flies. Multifactorial analysis revealed that AOX expression induces a significant change in only 0.8% of the 13,687 transcripts identified (Supplemental Table S16), affecting minor functional clusters (Supplemental Figure S6B), consistent with the idea that larval adaptation occurs mostly at the physiological level, with only a small impact on pupal development and survival. The LN diet affected 1.9% of all pupal transcripts (Supplemental Table S15), of which the major functional clusters were, once more, proteolysis and chitin metabolism, in addition to tissue integrity and transcriptional regulation (Supplemental Figure S6A). A much larger set of transcripts, 5.4% of the total, were altered by the LN-AOX interaction (Supplemental Table S17), the major affected clusters being stress response, transcriptional regulation, tissue integrity and stimulus response (Supplemental Figure S7), which may be markers of developmental failure. Because the LN and LN-AOX sets both included a mitochondrial cluster with transcripts for several OXPHOS subunits (Supplemental Figures S6A and S7), we analyzed in more details the transcripts coding for all subunits of the OXPHOS complexes, using manual clustering. We observed a general upregulation of OXPHOS transcripts in the LN-cultured, control flies (Figure 5A), which may reflect a response to maintain ATP production under conditions of lower nutrient availability.

**Figure 5.**
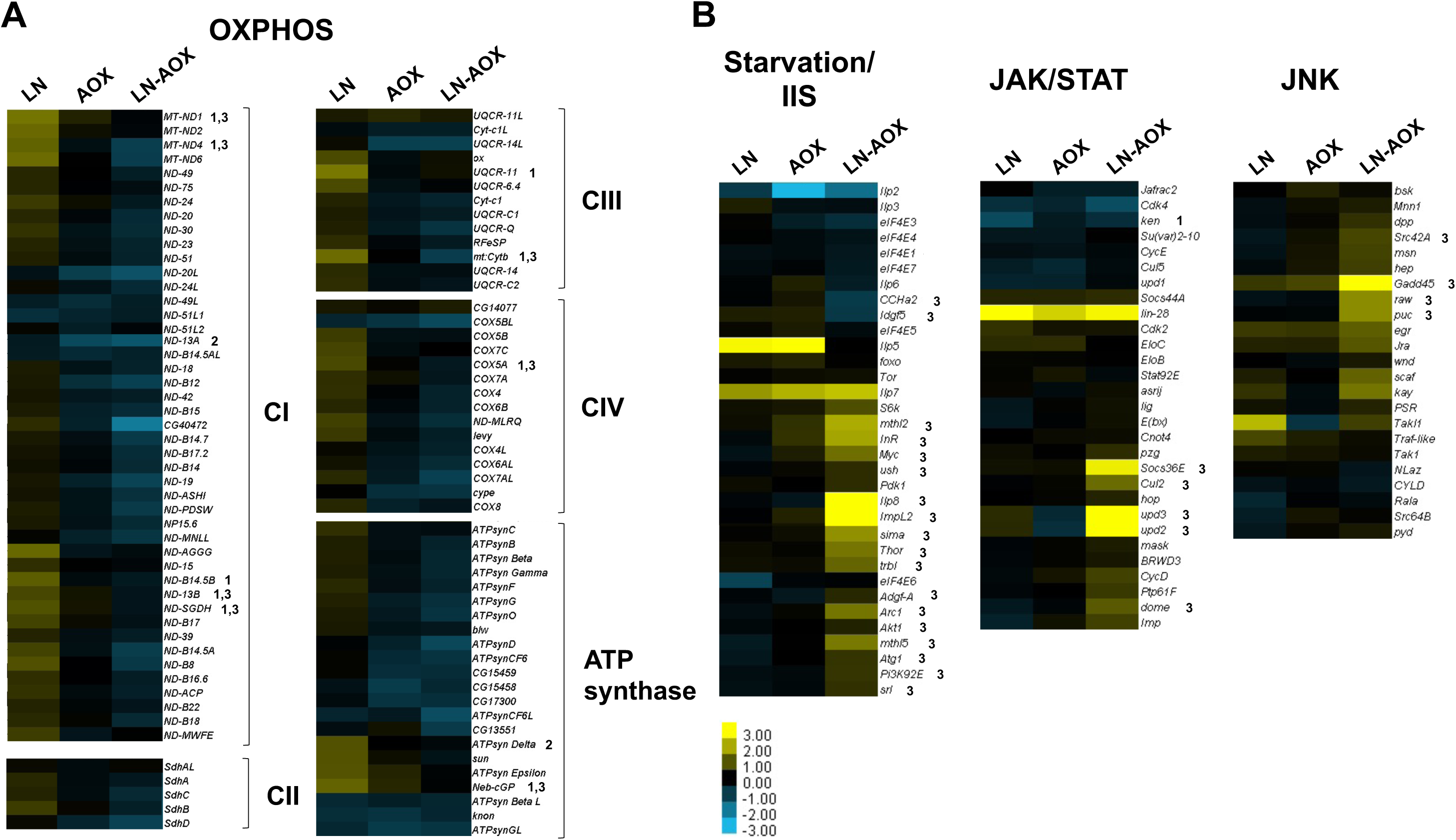
The interaction between strong ubiquitous expression of AOX and low nutrient (LN) diet promotes OXPHOS downregulation and a transcriptional response indicative of starvation, at the pupal stage. Heatmaps show changes in the levels of pupal transcripts encoding OXPHOS subunits (A) and of transcripts related to starvation response and signaling pathways as indicated (B). CI-IV, complex I-IV. 1-3 respectively indicate transcripts significantly (p < 0.05) altered by the LN diet, AOX expression and the LN-AOX interaction. Other details as in Figure 2.

Conversely, AOX induced a downregulation of OXPHOS transcripts in both diets, with the effect more pronounced in LN-AOX pupae (Figure 5A), which also showed a ∼12 fold increase in *lactate dehydrogenase* (*Ldh*) mRNA (Supplemental Table S17). High Ldh and low OXPHOS levels suggest that the LN-AOX interaction reflects mitochondrial dysfunction or failure of a metabolic switch during a developmental phase that relies on oxidative metabolism.

Upregulation of stress responses in pre-pharate LN-AOX pupae (Supplemental Figure S7) may indicate nutrient depletion, and that OXPHOS downregulation and other transcriptomic changes are downstream consequences of starvation. Accordingly, the levels of *Insulin-like receptor* (*InR*) transcripts were ∼3.2 fold higher in LN-AOX pupae. Moreover, manual clustering of transcripts associated with starvation-mediated response and Insulin/Insulin-like growth factor-1 Signaling (IIS) indicated a clear upregulation of these pathways in LN-AOX pupae (Figure 5B). For example, levels of *ImpL2*, a homolog of the human insulin-like growth factor binding protein-7 (IGFBP-7) gene, whose product is a secreted factor that binds the *Drosophila* insulin-like peptide (Ilp) 2 and inhibits growth upon nutritional deprivation (43), was elevated >10 fold. Another growth inhibitor highly expressed in LN-AOX pupae is *Ilp8* (∼21 fold higher); its encoded peptide is produced by the imaginal discs to delay pupariation in larvae (44, 45) and to arrest metamorphosis in early pupae (46). In both cases, Ilp8 inhibits ecdysone biosynthesis by the prothoracic gland when the imaginal discs show abnormal developmental growth. Interestingly, *Ilp8* transcripts already showed a tendency towards elevation (∼2.3 fold) in the LN-AOX larvae (Figure S8).

Upregulation of *Ilp8* in imaginal discs appears to be controlled by the Jun N-terminal Kinase (JNK) signaling pathway (44–46), which can be activated upon loss of tissue integrity(47). JNK signaling activates the JAnus Kinase/Signal Transducer and Activator of Transcription (JAK/STAT) cell-proliferating cascades in diverse tissues via upregulation of the cytokines Upd, Upd2 and Upd3 (47–50). We have shown previously that during metamorphosis AOX expression in the fly thoracic dorsal epithelium rescues engineered downregulation of the JNK pathway (27). Accordingly, we observed that LN-AOX pupae showed a general upregulation of JNK and JAK/STAT signaling. Particularly, the transcript levels of *upd2*, *upd3*, and their receptor *dome* were elevated ∼24, ∼46 and 2.3 fold, respectively (Figure 5B). Interestingly, diverse cell signaling-related genes are represented in the largest cluster of altered transcripts in the LN-AOX larval transcriptome (Supplemental Figure S3). In addition, AOX expression caused a significant increase in the larval levels of *upd2* and *upd3* independently of diet (Supplemental Figure S8), but their levels were normal in SD-cultured pupae (Figure 5B), whereas they increased further on LN diet. This might suggest that JAK/STAT signaling has a role in the larval physiological adaptation to AOX in SD, which may involve elevated fat storage, enabling the pupae to adjust to decreased bioenergetic efficiency yet still complete development. On LN diet, JAK/STAT signaling may not be effective in promoting this adaptation due to the reduced nutrient availability, leading to low larval fat storage. We thus propose that most LN-AOX pupae run out of metabolic fuel during metamorphosis, with associated changes in IIS, JNK and JAK/STAT signaling.

### Partial rescue of AOX-induced pupal lethality by methionine and/or tryptophan supplementation

Because the changes in larval metabolism appear to be key to our understanding of this pupal lethality, we also investigated the whole-body metabolome of AOX-expressing and control larvae cultured on SD and LN diets. A multifactorial analysis revealed that the LN diet causes significant alterations in the levels of 55% of the 74 unique metabolites identified (Supplemental Table S18). As expected, the LN diet promoted decreased levels of most intermediates of the TCA cycle and the pentose phosphate pathway, most amino acids, and a general rearrangement in the levels of nitrogenous bases, nucleosides and nucleotides, including an increase in markers of energy depletion such as nucleotide monophosphates and NAD+ (Figure 6). These changes are consistent with a combination of high energy demand for biosynthesis and storage of new biomolecules (characteristic of the proliferative nature of larval tissues) and low supply of high energy nutrients from the diet.

**Figure 6.**
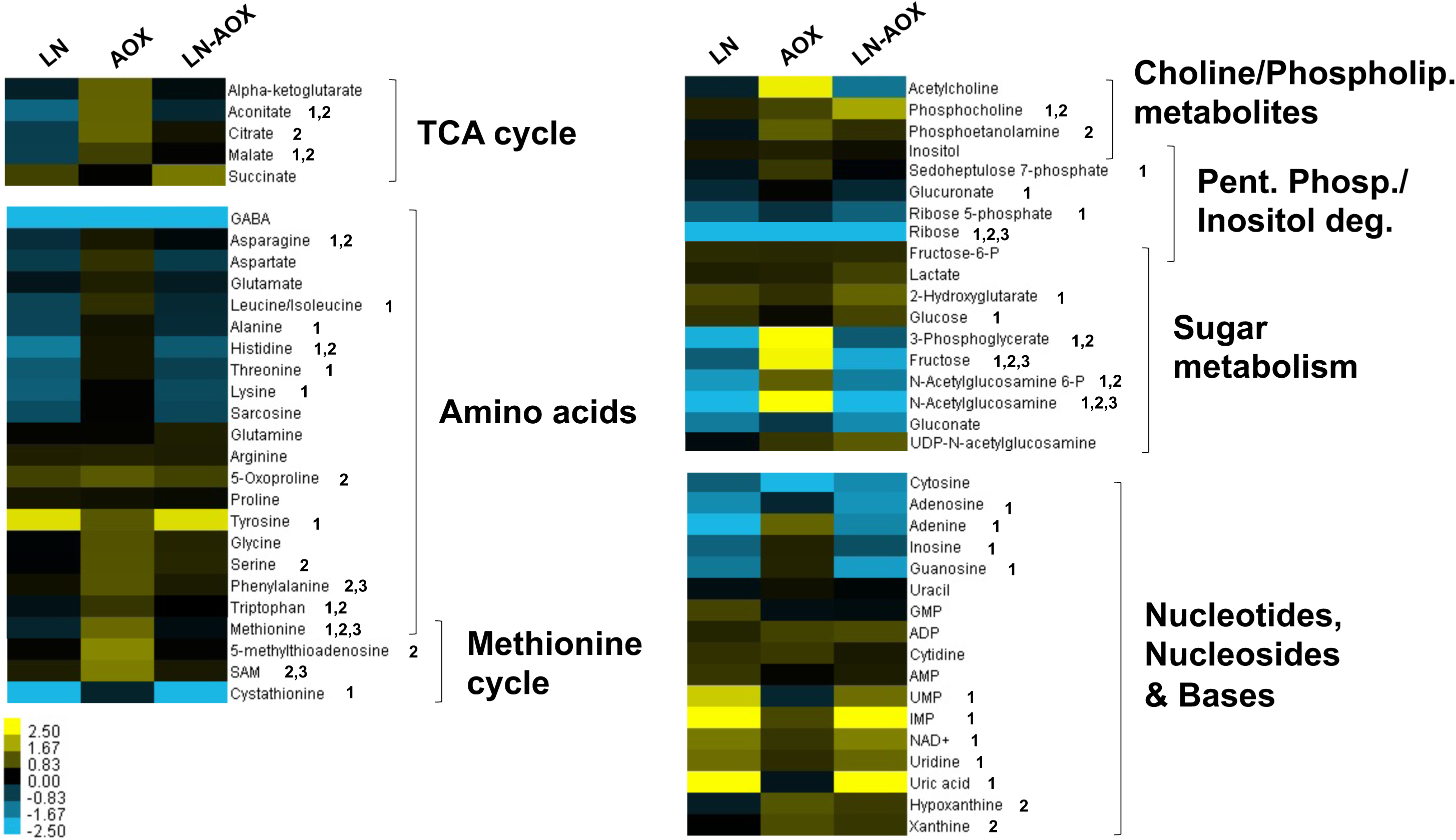
Larval amino acid levels are differentially affected by diet and AOX expression. Log2(fold change) values of the indicated metabolite in control larvae cultured on LN diet, AOX-expressing larvae cultured in standard (SD) diet, and AOX-expressing larvae culture in LN diet, each in a pairwise comparison with control larvae cultured in SD diet. 1-3 respectively denote metabolites significantly altered by the LN diet, AOX expression and the LN-AOX interaction (two-way ANOVA, p < 0.05). See Supplemental Table S18 for full list of metabolites. TCA, tricarboxilic acid; Phospholip., phospholipid; Pent. Phos., pentose phosphate pathway; Inositol deg., inositol degradation pathway. Control larvae, progeny of *w^1118^* and *daGAL4*; AOX-expressing larvae, progeny of *UAS-AOX^F6^* and *daGAL4*.

AOX expression, on the other hand, led to increased levels of most TCA cycle intermediates (Figure 6), suggesting important modifications of mitochondrial metabolism that were not evident from the larval transcriptomes. Succinate levels were not elevated though, which indicates that it (and/or succinyl-CoA, which was not detected in our analysis) might be diverted from the TCA cycle. We speculate succinate/succinyl-CoA may have been fueling the methionine cycle, of which several metabolites were elevated due to AOX expression (Figure 6). The methionine cycle is important in the cellular production of the universal methyl donor *S*-adenosylmethionine (SAM), but most importantly, it is interconnected with the metabolism of several other amino acids and nucleotides needed during tissue proliferation (51, 52). The amino acids serine, glycine and tryptophan, among others, can fuel the folate cycle, which in turn provides methyl groups via 5-methyl-tetrahydrofolate (5-methyl-THF) to the methionine cycle; together, the two cycles are referred to as one-carbon metabolism (53–55). Tryptophan, serine, glycine and a precursor in the serine *de novo* biosynthesis pathway, 3-phosphoglycerate (3-PG, possibly diverted from glycolysis/gluconeogenesis), were also elevated due to AOX expression, although the data for glycine were non-significant (Figure 6). Interestingly, whereas AOX expression tend to increase or preserve amino acid levels, methionine and phenylalanine were the only amino acids significantly altered by AOX in a diet-dependent manner (Figure 6).

We previously attempted to provide extra dietary protein by increasing the concentration of yeast extract up to 10% (w/w) into the LN diet, and observed no improvement in pupal viability upon AOX expression (22). Although rich in proteins, yeast extract generally contains proportionally low levels of the essential amino acids methionine and tryptophan, as well as the non-essential cysteine (56). Accordingly, the larval metabolome showed decreased levels of methionine and tryptophan (but not phenylalanine) due to LN diet. AOX expression, on the other hand, caused a significant increase in the levels of these amino acids, as well as of serine, glycine, and 3-PG, but this increase was either abolished or diminished in the metabolome of LN-AOX larvae (Figure 6). Methionine and tryptophan, among other essential (arginine and valine) and non-essential (glutamine, cysteine and aspartate) amino acids, were enriched in the water-soluble fraction of molasses that, when added to the LN diet, rescued the lethal phenotype caused by the LN-AOX interaction (22).

We therefore tested the supplementation of the SD and LN diets with methionine and tryptophan, individually and in combination, and observed a significant, albeit partial, rescue of lethality (Figure 7A) and of body size (Supplemental Figure S4B-E) of AOX flies cultured on LN diet. The effects were most apparent with low doses of tryptophan and were not additive, implying that a correct balance of these molecules is required for development, and suggesting that they are contributing to the same mechanism. In contrast, supplementing the LN diet with proline and/or glutamate failed to rescue the AOX-induced lethality (Figure 7B). We previously found that addition of extra sugar to the LN diet did not improve the survival of LN-AOX flies (22). Since sugar, proline and glutamate oxidation would provide reducing power for ATP synthesis via OXPHOS, we propose that energy production *per se* is not a limiting factor for LN-AOX larval metabolism. Although all our fly lines are in the *w^1118^* background, which is limited in tryptophan transport (57), our numerous control crosses (Figure 1) produced flies with varying expression levels of the transporter encoded by *white* (used as marker for the original transgenesis events that created the lines) with no apparent effect on viability on LN diet, confirming that the lethality is specifically caused by AOX expression. These results are consistent with the idea that yeast extract alone has an imbalanced amino acid content which impairs proper development of AOX-expressing flies. Total rescue of the AOX-driven pupal lethality, as seen with the addition of molasses to the LN diet (22), might require a combination of proper levels of methionine or tryptophan, plus other nutrients/molecules, which are yet to be identified. Elucidating this may not be straightforward, as even on SD diet, pupal viability of AOX-expressing flies seemed sensitive to the highest dose of methionine supplementation tested (Figure 7A), which may indicate an optimal dose, above and below which lethality increases.

**Figure 7.**
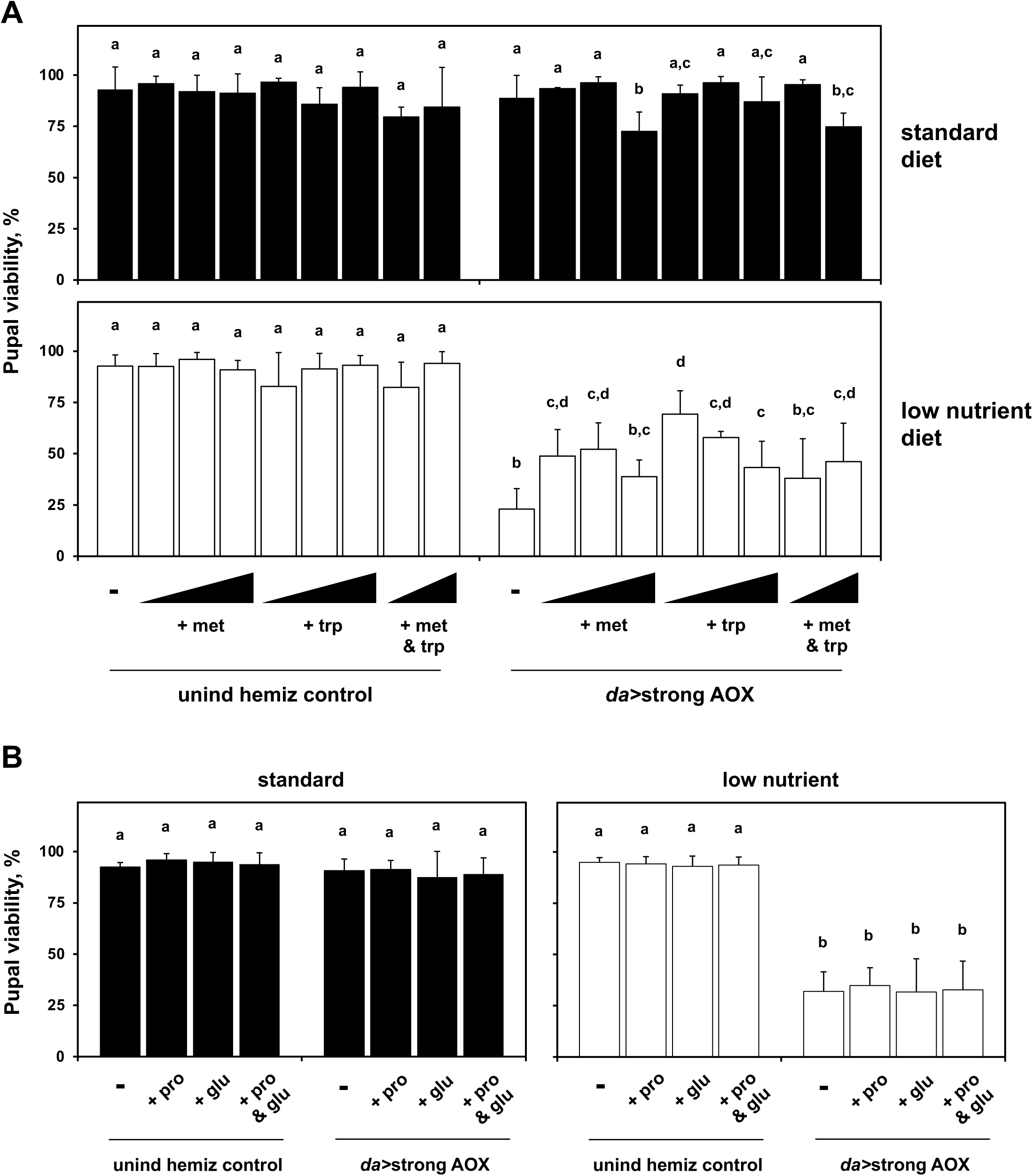
Partial rescue of AOX-induced pupal lethality on LN diet by methionine/tryptophan supplementation. Pupal viability (adult eclosion percentage) of flies of the indicated genotypes, cultured on the diet and supplements as shown. The final concentrations of added amino acids were: methionine (met) – 0.18, 0.35 and 0.7 mM; tryptophan (trp) – 0.1, 0.2 and 0.4 mM; proline (pro) and glutamate (glu) – 1.7 and 2.9 mM, respectively. These concentrations were selected based on previously published studies (96). unind hemiz controls, progeny of *UAS-AOX^F6^* and *w^1118^*; *da*>strong AOX, progeny of *UAS-AOX^F6^* and *daGAL4*. a-d represent significantly different statistical classes (one-way ANOVA, followed by the Tukey *post-hoc* test, p < 0.05).

The metabolomics analysis also revealed other metabolic pathways connected to the TCA cycle that might be of interest, in particular the γ-aminobutyric acid (GABA) shunt. The levels of GABA dropped very substantially (∼43 fold) in larvae cultured on LN diet (Figure 6 and Supplemental Table S18), suggesting it may have been catabolized into succinic semialdehyde and/or γ-hydroxybutyrate to bolster succinate levels. This would shift the equilibrium of the TCA cycle reaction catalyzed by succinate-CoA ligase towards production of succinyl-CoA (58), which we speculate is in contrast to the effects of AOX expression. Furthermore, the metabolism of chitin may also be an important target for further study: for all three conditions analyzed, the transcriptomics data indicate general rearrangements in the expression level of many chitin-related genes (Figure 2), and the metabolomic analyses show varying effects on the levels of N-acetylglucosamine-6-phosphate (Figure 6), an important intermediate for both catabolism and anabolism of chitin. More thorough investigations are warranted to address whether and how the interactions among the metabolic pathways mentioned above indeed play a role in the development of AOX-expressing flies.

### AOX-expressing larvae exhibit metabolic markers of enhanced growth

In our larval metabolomics analysis, we also found elevated levels of 2-hydroxyglutarate (2-HG) caused LN diet (Figure 6). Although our assay cannot distinguish between the L and D enantiomers, GABA metabolism inside mitochondria could increase the levels of D-2-HG (58). However, the *Drosophila* larva has been shown to produce substantial amounts of L-2-HG, derived from α-ketoglutarate by the action of Ldh in the cytoplasm (59, 60). AOX expression in the larvae causes upregulation of *Ldh* transcripts (Supplemental Table S10) and a trend towards accumulation of lactate, that was more pronounced under LN diet (Figure 6). This, in turn, may keep L-2-HG levels high by acidifying the cellular environment and inhibiting the breakdown of this metabolite (60, 61). Interestingly, both forms of 2-HG and succinate (Figure 6) are often considered oncometabolites (62), and are inhibitors of α-ketoglutarate-dependent protein and DNA demethylases (63). Their accumulation has independently been shown to cause epigenetic modifications (61,64–66), and may also cause post-translational modifications in key enzymes, which we speculate may drive important changes in gene expression and/or metabolic flux that are necessary for the *Drosophila* larva to adapt to AOX activity. The imbalanced levels of methionine/tryptophan (and likely other molecules) in the LN diet might actually be one of the mechanisms that impede these changes from occurring in AOX-expressing flies.

In our previous report (22), we showed that supplementation of the LN diet with a multivitamin complex containing folic acid did not altered the low viability rates of LN-AOX pupae, perhaps because conversion of folic acid to 5-methyl-THF is dependent on a source of methyl groups, such as serine or glycine. Their levels in the LN diet might be insufficient for proper development of AOX-expressing flies, but this has yet to be determined. Induction of the methionine cycle and production of SAM by methionine supplementation would bypass the need for other sources of methyl donors, but excess methionine can also compromise 5-methyl-THF synthesis by inhibiting the folate cycle enzyme methylene-THF reductase (67, 68), explaining the dose-dependent partial rescue of the AOX-induced lethality seen in Figure 7A. Synthesis of 5-methyl-THF using formate derived from tryptophan catabolism, on the other hand, would not cause such inhibition (68), perhaps explaining why LN supplementation with tryptophan alone in general increased pupal viability of LN-AOX flies better than supplementation with methionine alone, or methionine and tryptophan combined (Figure 7A).

Metabolomics analysis also suggests that the AOX-expressing larvae had altered choline metabolism (Figure 6). Oxidation of choline contributes to 5-methyl-THF synthesis and supplies electrons to the mitochondrial ETS (68), providing further support for the possible role of the one-carbon metabolism in the development of AOX-expressing larvae. Importantly, the changes in choline metabolites, which were apparent in the larval metabolomes, also suggest a link to the altered content of larval fat that we hypothesize above. The choline metabolite phosphocholine, whose levels are significantly affected by LN diet and AOX expression (Figure 6), is both a precursor and a catabolic product of the phospholipid phosphatidylcholine (41, 69). Phosphoethanolamine and inositol, whose levels trended higher due to AOX expression (Figure 6), are part of the polar head of the different forms of the phospholipids phosphatidylethanolamine and phosphatidylinositol. These phospholipids, along with the diversity of sterol types that can support *Drosophila* development (70), form the typical bilayer structure of cellular membranes and ultimately help control tissue integrity and function by regulating membrane integrity and selectivity (41, 71). Further experiments are needed to show the links relating one-carbon, amino acid, and phospholipid metabolism for the development of LN-AOX flies.

In addition to the inferred dependence on one-carbon metabolism for larval growth and completion of metamorphosis, AOX-expressing larvae showed many metabolic markers of increased tissue growth. These include: 1) elevated *Ldh* levels (Supplemental Table S10), associated with a trend to accumulate lactate and 2-HG (Figure 6); 2) accumulation of glycolytic and TCA cycle intermediates (Figure 6) that could be used in the biosynthesis of amino acids, lipids, and nucleotides, among other molecules; 3) dependence on an extra source of essential amino acids, such as methionine and/or tryptophan (Figure 7); 4) abnormal choline and membrane phospholipid metabolism (Figure 6 and Supplemental Figure S5); 5) altered signaling pathways (Figure 5 and Supplemental Figure S8); among others (52,55,69,72,73). Furthermore, proliferative tissues, including most tumor types, need mitochondria with a functional ETS, so that NADH and CoQH_2_ can be re-oxidized, allowing continuing or increased metabolic flux to meet the needs of biosynthesis (2). It has been shown that AOX expression is able to restore the overproliferative nature of cancer cells that are defective in CIII function and to promote tumors in mouse models (2). It is therefore possible that the expression of AOX in rapidly growing tissues/systems that contain functional CIII and CIV, such as *Drosophila* larvae, may increase metabolic flux by increasing NADH and CoQH_2_ reoxidation.

In this hypothetical scenario, the reactions of the TCA cycle that promote mitochondrial NAD+ regeneration for CI function would be accelerated, which in turn could also drive cataplerosis and induce larval growth, consistent with the changes in metabolism summarized above, and with the proposed increase in intestinal nutrient absorption and larval food consumption upon AOX expression (Figure 3). However, one would also expect increased larval biomass accumulation and/or faster development by AOX-expressing flies, at least when cultured on SD diet, but these have not been observed. On the contrary, we report here that AOX expression restricts larval weight gain even on SD diet, reaching ∼40% of biomass loss on LN diet, when compared with controls cultured on SD diet (Figure 4). We hypothesize that the limitations on biomass accumulation might be caused specifically by the strong expression of AOX (as is the high rate of pupal lethality), which in addition to increased cataplerosis, may also lead to effective mitochondrial uncoupling, with concomitant production of excess heat and/or compromise of mitochondrial ATP synthesis. Mitochondrial uncoupling and heat production by AOX has been previously evidenced in cultured human cells (6, 74). We next tested whether lower levels of AOX expression, which we showed not to cause lethality in combination with LN diet (Figure 1D), would have a positive impact on larval weight gain. In this case, the extent of mitochondrial uncoupling would not be as high (and ATP synthesis would perhaps be sufficiently compensated by substrate-level phosphorylation), but there would still be an increase in TCA cycle cataplerosis due to increased NADH and CoQH_2_ reoxidation, which could drive tissue growth. We thus selected the fly line 3x*tubAOX* (3), for which we have strong evidence of AOX-induced thermogenesis (and thus mitochondrial uncoupling) (21, 75), and in which AOX is ubiquitously and constitutively expressed, but at least 5 fold less (20) than in the flies with strong (GAL4-driven) expression of AOX used throughout this work. The AOX expression level in the 3x*tubAOX* flies is equivalent to what we denote as “intermediate” in Figure 1, which does not alter pupal viability, regardless of diet. We observed a significant gain of 10-15% in larval size, and in wet and dry masses when these flies were cultured on SD diet (Figure 8), which is in accordance with the idea that AOX expression induces growth in a dose-dependent fashion.

**Figure 8.**
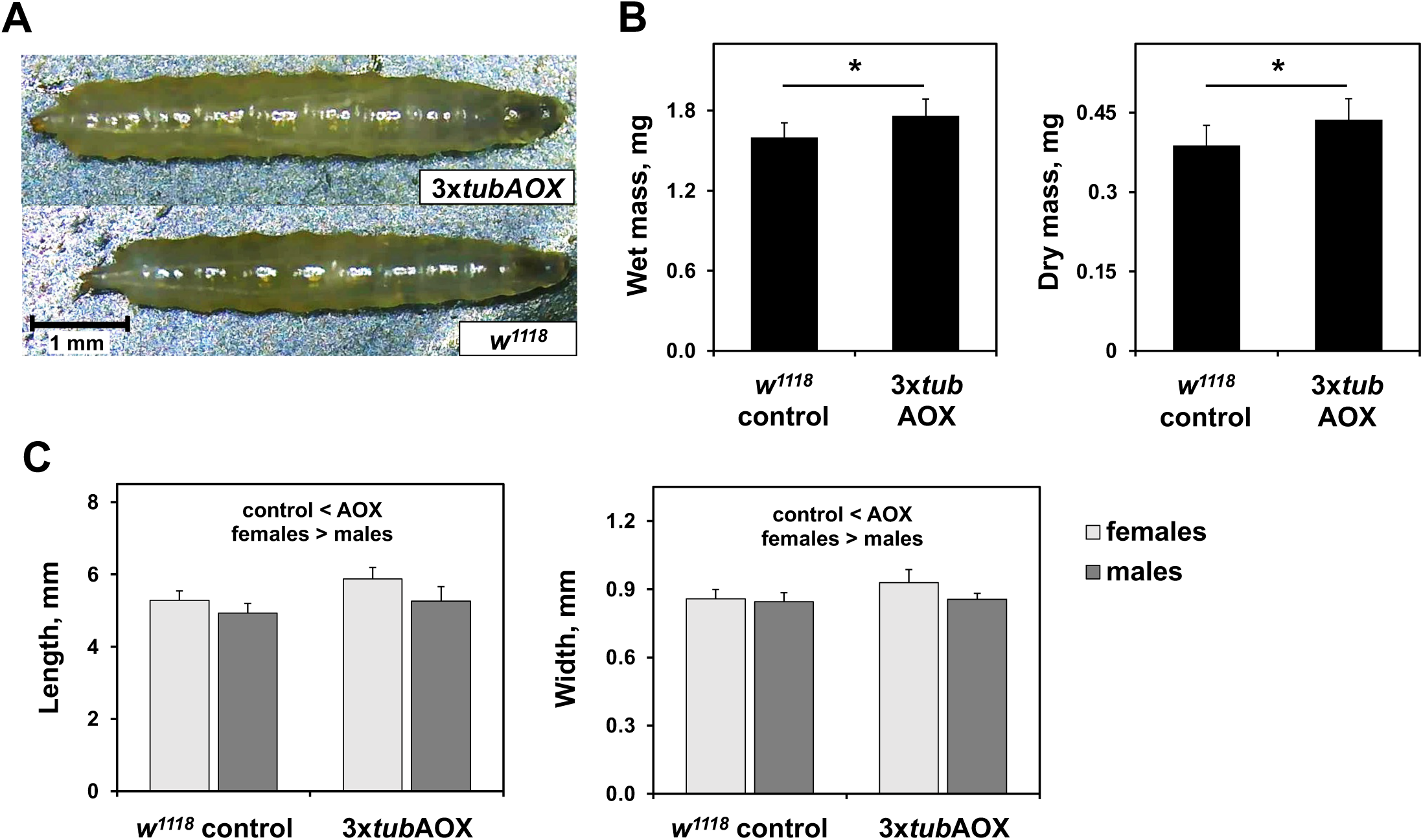
Modest levels of ubiquitous AOX expression promote larval growth. 3X*tubAOX* (3) larvae cultured on standard diet visualized by light microscopy (A, representative specimens), measured for wet and dry mass (B) and for dimensions as indicated (C). Values are averages from at least 25 individual specimens; error bars represent standard deviations. * indicates significant differences (Student’s t-test, p < 0.001). “control < AOX” indicate significant differences between control and AOX-expressing flies; “females > males”, between sexes (two-way ANOVA, p < 0.05).

These results, in combination with all other pieces of evidence we present here, suggest that tissue growth and biomass accumulation can be modulated by the xenotopic expression of AOX, the outcome of which is determined by the balance between increased metabolic flux, increased mitochondrial uncoupling, and diet.

### Conclusions

OXPHOS bypass using AOX has been proposed as a potential strategy for future treatments of human mitochondrial and related diseases. To validate its safety, work with AOX-expressing animal models is crucial. We confirm here that the development of flies with strong ubiquitous expression of AOX is significantly impacted when cultured on LN diet. In L3 larvae, transcriptomic changes are consistent with changes in intestinal morphology and in feeding behavior, suggesting that AOX induces (directly or indirectly) an increased need for dietary nutrients. AOX-expressing larvae also exhibit metabolic markers of enhanced growth, and higher storage of proteins and lipids. However, LN diet drastically inhibits biomass accumulation in the form of lipids by AOX-expressing larvae, likely resulting in disturbed signaling processes, increased stress and starvation responses, and ultimately, developmental arrest at the pupal stage. This arrest was partially rescued by LN supplementation with methionine or tryptophan, indicating a possible role for one-carbon metabolism in the successful development of AOX-expressing flies. Developmental arrest is dependent on the level and/or tissue pattern of AOX expression. In fact, lower (intermediate) ubiquitously-driven AOX-expressing larvae accumulate more biomass on SD diet, and have significantly better developmental outcome under cold stress (21), suggesting enhanced growth, even though energy dissipation as heat might still occur because of mitochondrial uncoupling. Taken together, our work suggests that AOX may hormetically modulate growth by a balance between increased metabolic flux and increased mitochondrial uncoupling: the former promoting TCA cycle cataplerosis and anabolism, to a point at which the latter starts to affect ATP synthesis and/or produce too much metabolic heat, ultimately restricting growth (Figure 9). It is also possible that the antioxidant effects of AOX (5, 6) interfere with ROS signaling, which are essential mediators of differentiation, growth and stress (76–79), but this has yet to be determined.

**Figure 9.**
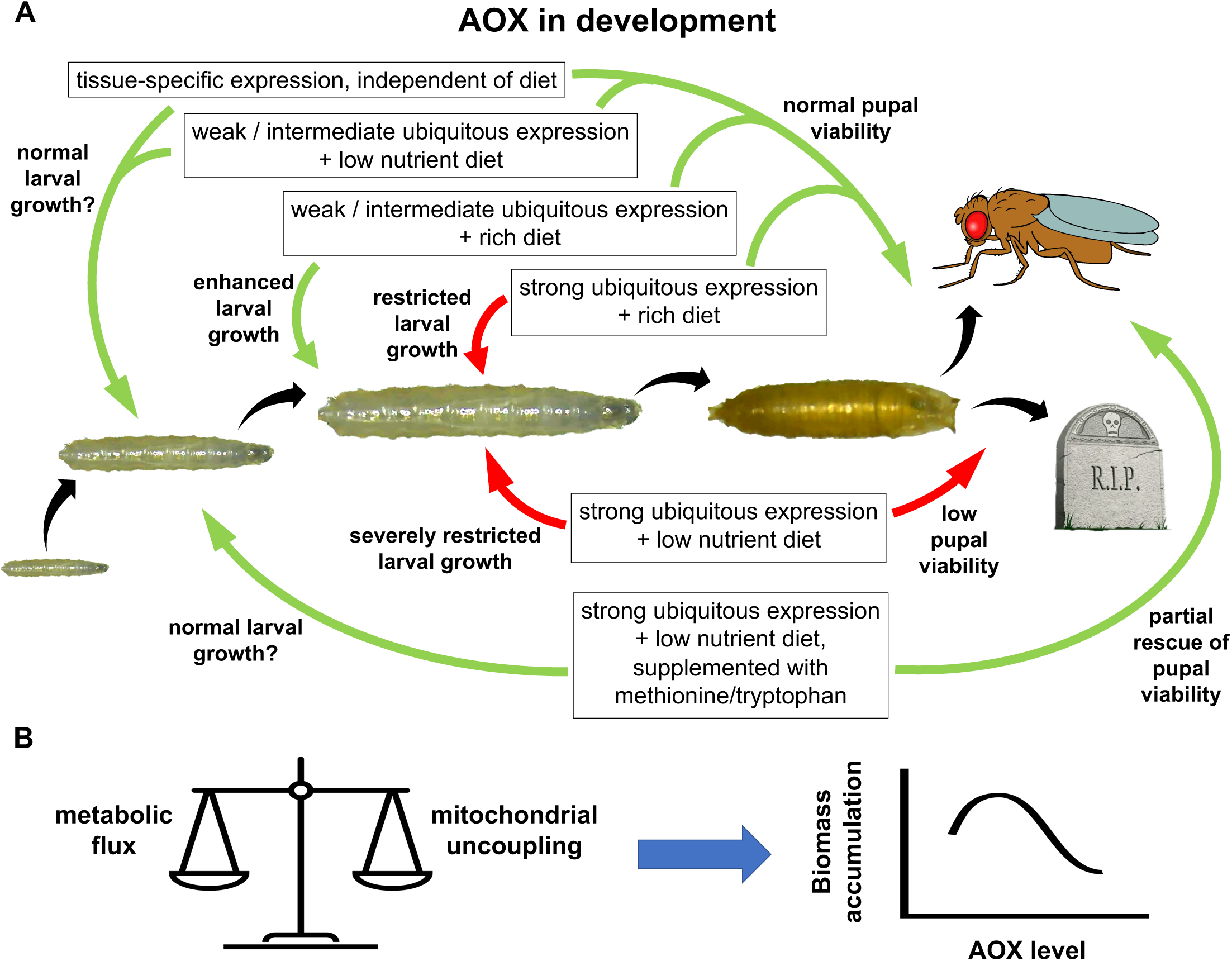
AOX expression during *Drosophila* development. A, Summary of the results presented in this work. B, Proposed model of how AOX influences larval growth and biomass accumulation, impacting pupal viability and adult eclosion rates. See text for details.

Because the developmental arrest described here is likely dependent on a systemic impact caused by strong ubiquitous expression of AOX in combination with LN diet, the metabolic and signaling changes induced by AOX possibly depend on the functions of a particular tissue or organ. AOX expression might influence general larval growth by acting locally in each tissue/organ, but the problems seen on LN diet occur with the integration of these organs’ functions. The musculature and the fat body, for example, are supposedly responsible for the storage of the extra protein and fat in AOX-expressing larvae, respectively, and being the tissues with the most elevated biosynthetic need for nutrients, their mitochondria would participate significantly in the respective anabolic processes. However, to supply the extra precursor molecules needed, the nervous system and the intestines would need to be more active in inducing increased larval feeding behavior, in digesting the extra food consumed, and in absorbing the extra nutrients. Mitochondria in these tissues would be primarily catabolic, with a high need to synthesize ATP. By applying whole-larva omics, we may have missed important tissue-specific signals and metabolic changes promoted by AOX. Therefore, future studies should focus on profiling tissue-specific metabolic disturbances caused by AOX in the growing *Drosophila* larva, and how they are integrated to promote growth and biomass accumulation (or the lack thereof in LN diet).

The developing *Drosophila* larva has already been postulated as a model for our understanding of the metabolism of animal tissue growth, including that of cancer. It might even be a more realistic model than cancer cells cultured *in vitro*, since one can study how a complex living organism regulates metabolic flux intrinsically and in peripheral tissues, supporting the transitions through distinct developmental stages with varying metabolic demands (80). The idea that certain levels of AOX expression promote growth is worrisome from the point-of-view of putative therapeutic use, given that it remains unclear whether the effects are merely hyperplastic, hypertrophic or also neoplastic. However, although highly speculative, the fact that the combination of strong expression of AOX and poor diet severely restricts growth may also indicate their potential applicability in cancer research. Better understanding the dose- and diet-dependent effects of AOX in tissue growth in the *Drosophila* larva will, therefore, be extremely relevant.

## Materials and Methods

### Fly stock and diets

The *D. melanogaster* recipient line *w^1118^*, the driver lines *daughterless-GAL4* (*daGAL4*) (81), *embryonic lethal abnormal visionC155-GAL4* (*elavGAL4*) (82), *myosin heavy chain-GAL4* (*mhcGAL4*) (83) and *r-tetramer-GAL4* (*r4GAL4*) (84), and the GAL4-dependent *UAS-GFP^Stinger^* line (85), were obtained from stock centers. The driver line *drumstick-GAL4* (*drmGAL4*) (38) was a gift from Dr. Lucas Anhezinís lab. Previously described transgenic lines carrying *tubAOX*, *UAS-AOX* constructs or the *UAS* promoter only (“empty” pUASTattB plasmid) were used: 3x*tubAOX* (inserted into chromosomes X, 2 and 3)(3), *UAS-AOX^F6^* (inserted into chromosome 2) (10), *UAS-AOX^8.1^* (chromosome 2), *UAS-AOX^7.1^* (chromosome 3), *UAS-mutAOX^2nd^* (chromosome 2), *UAS-mutAOX^3rd^* (chromosome 3) and *UAS-empty^2nd^* (chromosome 2) (16). The double transgenic lines *UAS-AOX^8.1^;UAS-AOX^7.1^* and *UAS-mutAOX^2nd^;UAS-mutAOX^3rd^* were created by standard genetic crosses using lines carrying traditional balancer chromosomes. All fly lines were backcrossed with the recipient line *w^1118^* six-to-ten generations prior to the experiments. The fly lines were maintained on SD diet (10), with a 12/12h of light/dark photoperiod cycle at 18 or 25 °C. The LN diet consisted of 3.5% yeast extract, 1% agar and antimicrobials (0.1% niapigin and 0.5% propionic acid). Where indicated in figures and legends, diets were supplemented with the amino acids methionine, tryptophan, proline and glutamate at the concentrations shown.

### Developmental assays

The *UAS-AOX*, *UAS-GFP^Stinger^* and *UAS-empty* lines were crossed to either a *GAL4* driver line to induce transgene expression, or to the recipient line *w^1118^* (generating uninduced hemizygous controls), as indicated in the figure legends. Based on immunoblot analyses (Figure 1B) and previously published work (16), the crosses between *UAS-AOX^F6^* and the *GAL4* drivers generated flies with high expression levels of AOX, whereas the same crosses using *UAS-AOX^8.1^* or *UAS-AOX^7.1^* produced low levels of AOX. The double transgenic lines *UAS-AOX^8.1^;UAS-AOX^7.1^* and *UAS-mutAOX^2nd^;UAS-mutAOX^3rd^*, upon crossing with *GAL4* drivers, and the triple transgenic line 3x*tubAOX* (constitutive expression) generated flies with intermediate expression levels of AOX. Ten-fifteen female virgins and 5-7 males were pre-mated on SD diet during 24-48 h at 25 °C with a 12/12 h light/dark photoperiod cycle, flipped into vials containing SD or LN diets, and allowed to lay eggs until the limiting rate of 60-80 eggs per vial was reached. The eggs were allowed to develop in the same vials, from which the numbers of pupae and eclosing adults were recorded. Pupal viability was calculated as the ratio between eclosed adults and total number of pupae, averaged among 2-3 biological replicates, each carrying 5-8 vials (technical replicates).

### Measurements of body mass, intestinal length, food consumption, and body composition

For larval body mass measurements, batches of ten randomly selected, wandering L3 larvae (100-115 h post-egg laying in SD diet; 125-140 h in LN diet) were collected, briefly rinsed in deionized water, carefully dried on Kimwipes® sheets, placed in a pre-weighed 1.5 mL microcentrifuge tube, and weighed using a precision balance (ATX224, Shimadzu Scientific Instruments) to obtain the wet mass. The tubes were then frozen at -20 °C for up to 14 days, and placed in a dry bath at 65 °C with the lids open until the larval weight dropped to constant values (∼2 h), which were recorded as the dry mass. Next, the samples were incubated with 1 mL ethyl ether for 24 h at room temperature, and the procedure was further repeated three times, after which the samples were dried as described above, to obtain the lean mass. Individual larval masses were calculated by dividing the total weight per tube by 10, and the final values were obtained by averaging the individual larval masses of ∼40 tubes, obtained in three independent experiments. Measurements of total fat content were obtained by subtracting lean from dry mass.

For estimates of total protein content, batches of 20 randomly selected, wandering L3 larvae (100-115 h post-egg laying in SD diet; 125-140 h in LN diet) were collected, briefly rinsed in deionized water, carefully dried on Kimwipes® sheets, and homogenized on ice by 3-5 strokes of a hand-held homogenizer in 1 mL isolation buffer (250 mM sucrose, 5 mM Tris-HCl, 2 mM EGTA, pH 7.5). Total protein content was estimated by the Bradford method. Individual larval protein content was calculated by dividing the total protein per tube by 20, and the final values were obtained by averaging data from 4-6 independent experiments.

For measurements of intestinal length, randomly selected, wandering L3 larvae (100-115 h post-egg laying in SD diet; 125-140 h in LN diet) of each sex were collected, briefly rinsed in deionized water, and submerged in ∼80 °C water for ∼2 s for immobilization. The specimens were immediately photographed and then transferred to a dish containing PBS (8 mM Na_2_HPO_4_, 2 mM KH_2_PO_4_, 150 mM NaCl, 30 mM KCl, pH 7.4). The intestines were immediately dissected and photographed. The images were analyzed and intestinal length estimated using ImageJ v1.53e software. The ratio between intestinal length and larval area was averaged from measurements of ∼15 individuals of each sex, in three independent experiments.

Measurements of larval food consumption were adapted from (86). Briefly, randomly selected, early L3 larvae (75-85 h post-egg laying in SD diet; 95-105 h in LN diet) were collected, rinsed in deionized water, and placed on a dish without food for 20 min at 25 °C. The individuals were then transferred to a plastic petri dish with 6 mL of food containing 2 g yeast extract and 15 mg Brilliant blue R dye, and allowed to feed for 10 min, after which they were processed and their intestines were dissected and photographed, as described above. The images were analyzed and the blue signal intensity was estimated using ImageJ v1.53e software. The ratio between the blue signal intensity and intestinal length was averaged from measurements of 7-10 specimens, from at least two independent experiments.

### Immunoblotting

Mitochondrial protein extracts were prepared on ice by homogenization of 20 randomly selected, whole adult males (1-3 days after eclosion) of the indicated genotype in isolation buffer (250 mM sucrose, 5 mM Tris-HCl, 2 mM EGTA, pH 7.5), followed by centrifugation at 200 x *g_max_* for 1 min at 4 °C. The supernatant was removed and centrifuged again at 200 x *g_max_* for 3 min at 4 °C. The recovered supernatant was further centrifuged at 9,000 x *g_max_* for 10 min at 4 °C, and the mitochondrial pellet was resuspended in 50 μL isolation buffer. The protein quantification was performed by the Bradford method, and the samples were stored at -80°C until use. Forty μg of mitochondrial protein extract were mixed with Laemmli buffer (2% SDS, 10% glycerol, 5% 2-mercaptoethanol, 0.02% bromophenol blue and 62.5 mM Tris-HCl, pH 6.8), denatured at 95 °C for 3 min, and resolved on 12% SDS-polyacrylamide gels for ∼3.5 h at 80 V, alongside the Bio-Rad Precision Plus Protein Standards marker. The proteins were transferred at 550 mA for 7 min to nitrocellulose membranes using the semi-dry transfer system MSD10 (Major Science) at room temperature. The membranes were blocked using 5% nonfat milk/PBST solution (8 mM Na_2_HPO_4_, 2 mM KH_2_PO_4_, 150 mM NaCl, 30 mM KCl, 0.05% Tween 20, 5% dried nonfat milk, pH 7.4) for at least 2 h, and incubated with a 1% nonfat milk/PBST solution containing the primary antibodies anti-AOX (rabbit polyclonal, 1:20,000) (10) and anti-ATP5A (mouse monoclonal, Abcam, UK, 1:10,000) for 3 h at room temperature under constant shaking. After three 40 min washes in PBST, the membranes were incubated with a 1% nonfat milk/PBST solution containing the secondary HRP-conjugated goat anti-rabbit (1:10,000, Bio-Rad, USA) and anti-mouse (1:10,000, Bio-Rad, USA) antibodies overnight at 4 °C, and washed as described. Chemiluminescence signals were detected on a ChemiDoc Imaging System (Bio-Rad), after the membranes were incubated with the luminol substrate Immun-Star HRP (Bio-Rad).

### RNA extraction, sequencing and data analyses

Ten randomly selected, wandering L3 larvae (100-115 h post-egg laying in SD diet; 125-140 h in LN diet) and 10 pre-pharate pupae (65-80 h post-pupariation) per biological replicate (3 in total) were collected, briefly rinsed in ultrapure water to remove dietary residues, and immediately processed. Total RNA was extracted using the RNeasy® Tissue kit (QIAGEN), according to manufacturer instructions, eluted in nuclease-free water, and analyzed using the Qubit® RNA HS Assay kit (Invitrogen) and the RNA 6000 Nano LabChip kit/Bioanalyzer 2100 (Agilent Technologies). One µg samples of RNA with high integrity number (RINe > 8.9) were used for RNA sequencing library preparation using the TruSeq Stranded mRNA Library Prep Kit (Illumina), according to the manufacturer’s protocol. The libraries were quantified using the Promega QuantiFluor dsDNA System on a Quantus Fluorometer (Promega), and analyzed using the High Sensitivity D1000 Screen Tape on an Agilent 2200 TapeStation instrument. The libraries were normalized, pooled, and subjected to cluster generation, and paired-end sequencing was performed for 2x150 cycles on a HiSeq4000 instrument (Illumina), according to the manufacturer’s instructions, by the company Omega Bioservices (Norcross, GA, USA).

The sequencing reads were processed with Trimmomatic (87) and assessed by FastQC (88) for processing quality, following the steps available at (89). The downstream data analysis was performed through the Chipster v3.16 software package (90), according to the following steps. The alignment of the reads was performed with HISAT2 (91) using the *D. melanogaster* reference genome r6.30, available on FlyBase (34). The quantitation of mapped reads was performed by HTSeq (92); only reads with more than 5 counts in at least 3 libraries were considered for the analyses. The statistical analysis was performed using the edgeR v3.14 package (93) in the multifactorial or pairwise analysis mode, where indicated. The factors considered were diet, AOX expression, and the interaction between diet and AOX. Genes differentially expressed were determined as p < 0.05. Gene ontology analysis was performed considering each factor using DAVID v6.8 (94). Unsupervised clustering was performed by the Cluster3.0 software, and heatmaps were created through JAVA TreeView v1.1.6r4.

### Metabolite extraction and mass spectrometry analyses

Ten randomly selected, wandering L3 larvae (100-115 h post-egg laying in SD diet; 125-140 h in LN diet) per biological replicate (5 in total) were collected, briefly rinsed in ultrapure water, and homogenized in 1 mL cold (-80 °C) 80% methanol on dry ice using a probe homogenizer. Samples were clarified by centrifugation for 5 min at 16,000 x *g_max_* at 4 °C and the supernatants were transferred to new microcentrifuge tubes. The remaining pellets were resuspended in RIPA lysis buffer (10 mM Tris-HCl, 150 mM NaCl, 1% Triton X-100, 0.1% deoxycholate, 0.1% SDS, pH 7.5) and total protein content was determined by the BCA method. A volume of the supernatant equivalent to 5 µg of protein was transferred to a glass vial, dried using an EZ-2 Elite evaporator (SP Scientific), and stored at -80 °C until analysis by LC-MS. The samples were analyzed using an UltiMate 3000RSLC HPLC (Thermo Scientific) coupled to a Q Exactive mass spectrometer (Thermo Scientific). After resuspension in 50% acetonitrile, 10% of each sample was loaded onto a Luna NH2 (3 µm 100A, 150 mm x 2 mm, Phenomenex) column. Separation was achieved with 5 mM NH_4_OAc, pH 9.9 (mobile phase A) and ACN (mobile phase B) at a flow rate of 200 µL/min and a linear gradient from 15% to 90% A over 18 min. This was followed by an isocratic step at 90% A for 9 min and re-equilibration to the initial 15% A for 7 min. The Q Exactive was run with polarity switching (+3.50 kV / -3.50 kV) in full scan mode with an *m/z* range of 65-975.

Metabolites were identified with TraceFinder 4.1 (Thermo Fisher Scientific) using accurate mass measurements (≤ 3 ppm) and expected retention times previously established with pure standards. Quantities were determined by peak area integration and the statistical analysis was performed using the MetaboAnalyst5.0 software (95). Missing values were replaced by 1/5 of minimum positive values of the corresponding variables. Following the normalization step, a two-way ANOVA type I analysis was performed, including the interaction between the AOX and LN factors, and the metabolites significantly altered were determined as p-adjusted value < 0.05.

## Data Availability

Raw RNA-Seq and metabolomic data are deposited, respectively, in the NCBI Sequence Read Archive (BioProject PRJNA763657) and the NIH Common Fund’s National Metabolomics Data Repository (Project ID 3056). Processed data can be found in the Supplemental Tables 1-18. Raw data of the analyses presented in Figures 1, 3, 4, 7 and 8 can be obtained from the corresponding author upon request.

## Acknowledgements

We would like to thank Dr. Lucas Anhezini for providing the *drmGAL4* line, and Dr. Richard Hensh for help with the statistical analyses. A.F.C., M.M.C., G.S.G., M.F.O. and A.A.M. were supported by fellowships from the Coornadoria de Aperfeiçoamento de Pessoal de Nível Superior (CAPES, grant numbers 88882.434433/2019-01 and 88882.434441/2019-01), Conselho Nacional de Desenvolvimento Científico e Tecnológico (CNPq, grant number 141001/2019-4), and Fundação de Amparo à Pesquisa do Estado de São Paulo (FAPESP, grant numbers 2017/13743-2, 2020/07797-5 and 2017/17645-5). M.T.O. would like to acknowledge funding from FAPESP (grant numbers 2014/02253-6 and 2017/04372-0), CNPq (grant numbers 424562/2018-9 and 306974/2017-7) and the European Union (Marie Curie International Incoming Fellowship GA328988). E.D. acknowledges funding from AFM-Telethon (research grant number 17424) and the Academy of Finland (research fellowship number 130172). H.T.J. acknowledges funding from Academy of Finland (decisions 283157 and 307431) and the European Research Council (Advanced Grant 232738).

