## Supplemental Figures for "Systemic impact of the expression of the mitochondrial alternative oxidase on *Drosophila* development"

**Supplemental Figure Legends**

**Figure S1.** Altered transcriptional pattern in larvae triggered by the low nutrient (LN) diet. The heatmaps show the genes and gene clusters altered by LN diet, except the ones shown in Figure 2 (proteolysis and chitin metabolism). LN, AOX and LN-AOX show respectively color representations of the $\log_2$(fold change) values of the indicated transcript in control larvae cultured in LN diet, AOX-expressing larvae cultured in standard diet, and AOX-expressing larvae culture in LN diet, each in a pairwise comparison with control larvae cultured in standard diet. The indicated genes were selected as significantly differentially expressed ($p < 0.05$) by the LN diet, independent of AOX expression, according to a multifactorial analysis performed using RNA-seq data and the software edgeR[1]. DAVID functional annotation tool[2] was used to generate the indicated gene clusters (see Material and Methods for details). Prot. Fold., protein folding; RNA Bind., RNA binding; Glycol., glycolysis; Sex. Matur., sex maturation; Mitoc., mitochondrial; A.A. Transp., amino acid transport; Alk. Phos., alkaline phosphatase; F.A. Met., fatty acid metabolism; TM, transmembrane proteins. Control larvae, progeny of *UAS-AOX*$^{F6}$ and *w*$^{1118}$; AOX-expressing larvae, progeny of *UAS-AOX*$^{F6}$ and *daGAL4*.

**Figure S2.** Altered transcriptional pattern in larvae triggered by strong ubiquitous expression of AOX. The heatmaps show the genes and gene clusters altered by AOX, except the ones shown in Figure 2 (proteolysis and chitin metabolism). LN, AOX and LN-AOX show respectively color representations of the $\log_2$(fold change) values of the indicated transcript in control larvae cultured in low nutrient (LN) diet, AOX-expressing larvae cultured in standard diet, and AOX-expressing larvae culture in LN diet, each in a pairwise comparison with control larvae cultured in standard diet. The indicated genes were selected as significantly differentially expressed ($p < 0.05$) by AOX expression, independent of the LN diet, according to a multifactorial analysis performed using RNA-seq data and the software edgeR[1]. DAVID functional annotation tool[2] was used to generate the indicated gene clusters (see Material and Methods for details). Control larvae, progeny of *UAS-AOX*$^{F6}$ and $w^{1118}$; AOX-expressing larvae, progeny of *UAS-AOX*$^{F6}$ and *daGAL4*.

**Figure S3.** Altered transcriptional pattern in larvae triggered by the interaction between low nutrient (LN) diet and strong ubiquitous expression of AOX (LN-AOX). The heatmaps show the genes and gene clusters altered by the LN-AOX interaction, except the ones shown in Figure 2 (proteolysis and chitin metabolism). LN, AOX and LN-AOX show respectively color representations of the $\log_2$(fold change) values of the indicated transcript in control larvae cultured in LN diet, AOX-expressing larvae cultured in standard diet, and AOX-expressing larvae culture in LN diet, each in a pairwise comparison with control larvae cultured in standard diet. The indicated genes were selected as significantly differentially expressed ($p < 0.05$) by the LN-AOX interaction, according to a multifactorial analysis performed using RNA-seq data and the software edgeR[1]. DAVID functional annotation tool[2] was used to generate the indicated gene clusters (see Material and Methods for details). Detox., detoxification; Mitoc., mitochondrial; Memb. Prot., membrane proteins. Control larvae, progeny of *UAS-AOX*$^{F6}$ and $w^{1118}$; AOX-expressing larvae, progeny of *UAS-AOX*$^{F6}$ and *daGAL4*.

**Figure S4.** Changes in larval and pupal sizes caused by strong ubiquitous expression of AOX, and by dietary supplementation with methionine and/or tryptophan. A, quantitation of larval area of the indicated flies cultured on the indicated diet. The values indicate averages of 15 individual measurements, and the error bars represent standard deviations. B-C, light microscopy images of representative pupae: SD-control, control flies in standard diet; LN-control, control flies in low nutrient diet; SD-AOX, flies with strong expression of AOX in standard diet; LN-AOX, flies with strong expression of AOX in low nutrient diet; LN+met-AOX, flies with strong expression of AOX in low nutrient diet supplemented with 0.7 mM L-methionine; LN+trp-AOX, flies with strong expression of AOX in low nutrient diet supplemented with 0.4 mM L-tryptophan; LN+met+trp-AOX, flies with strong expression of AOX in low nutrient diet supplemented with both amino acids. D-E, quantitation of pupal length and width of the indicated flies cultured on the indicated diet. The values indicate averages of at least 7-10 individual measurements, and the error bars represent standard deviations. + Met, addition of 0.7 mM L-methionine; + Trp, addition of 0.4 mM tryptophan; + Met & Trp, addition of both amino acids; control, progeny of $UAS\text{-}AOX^{F6}$ and $w^{1118}$; AOX, progeny of $UAS\text{-}AOX^{F6}$ and *daGAL4*. a, b and c represent significantly different statistical classes, according to an one-way ANOVA, followed by the Tukey *post-hoc* test.

**Figure S5.** Changes in larval and pupal transcripts related to lipid metabolism. The heatmaps show selected genes whose products are involved with the indicated pathways. LN, AOX and LN-AOX show respectively color representations of the $\log_2$(fold change) values of the indicated transcript in control flies cultured on low nutrient (LN) diet, AOX-expressing flies cultured on standard diet, and AOX-expressing flies culture on LN diet, each in a pairwise comparison with control flies cultured on standard diet. Control flies, progeny of $UAS\text{-}AOX^{F6}$ and $w^{1118}$; AOX-expressing flies, progeny of $UAS\text{-}AOX^{F6}$ and *daGAL4*. 1-3 respectively indicate transcripts significantly ($p < 0.05$) altered by the LN diet, AOX expression and the LN-AOX interaction, according to a multifactorial analysis performed using RNA-seq data and the software edgeR[50].

**Figure S6.** Altered transcriptional pattern in pupae triggered independently by the low nutrient (LN) diet and strong ubiquitous expression of AOX. The heatmaps show the genes and gene clusters altered by LN diet (A) and AOX expression (B). LN, AOX and LN-AOX show respectively color representations of the $\log_2$(fold change) values of the indicated transcript in control pupae cultured on LN diet, AOX-expressing pupae cultured on standard diet, and AOX-expressing pupae culture on LN diet, each in a pairwise comparison with control pupae cultured on standard diet. The indicated genes in A were selected as significantly differentially expressed ($p < 0.05$) by the LN diet, independent of AOX expression, whereas the indicated genes in B were selected as significantly differentially expressed ($p < 0.05$) by AOX expression, independent of diet, according to a multifactorial analysis performed using RNA-seq data and the software edgeR[1]. DAVID functional annotation tool[2] was used to generate the indicated gene clusters (see Material and Methods for details). Cal. Reg. Proc., calcium regulated process; Mitoc., mitochondrial; Morpho, morphogenesis; Chit. Met., chitin metabolism; Tissue Integ., tissue integrity; Transc. Reg., transcriptional regulation; Proteol., proteolysis; Detox., detoxification; F.A. Met., fatty acid metabolism. Control pupae, progeny of *UAS-AOX$^{F6}$* and *$w^{1118}$*; AOX-expressing pupae, progeny of *UAS-AOX$^{F6}$* and *daGAL4*.

**Figure S7.** Altered transcriptional pattern in pupae triggered by the interaction between low nutrient (LN) diet and strong ubiquitous expression of AOX. The heatmaps show the genes and gene clusters selected as significantly differentially expressed ($p < 0.05$) by the LN-AOX interaction, according to a multifactorial analysis performed using RNA-seq data and the software edgeR[1]. DAVID functional annotation tool[2] was used to generate the indicated gene clusters (see Material and Methods for details). LN, AOX and LN-AOX show respectively color representations of the $log_2$(fold change) values of the indicated transcript in control pupae cultured on LN diet, AOX-expressing pupae cultured on standard diet, and AOX-expressing pupae culture on LN diet, each in a pairwise comparison with control pupae cultured in standard diet. Stress Resp., stress response; Proteol., proteolysis; Nerv. Sys., nervous system; Mitoc., mitochondrial; Chit. Met., chitin metabolism; Metal Reg. Proc., metal-regulated process; Tiss. Integ./Stimu. Resp., tissue integrity and stimulus response; Transc. Reg., transcriptional regulation. Control pupae, progeny of *UAS-AOX$^{F6}$* and *w$^{1118}$*; AOX-expressing pupae, progeny of *UAS-AOX$^{F6}$* and *daGAL4*.

**Figure S8.** Changes in larval transcripts related to signaling pathways. The heatmaps show changes in the level of larval transcripts related to starvation-mediated response and the Insulin/Insulin-like growth factor-1 Signaling (IIS) pathway, the JAnus Kinase/Signal Transducer and Activator of Transcription (JAK/STAT) signaling pathway, and the Jun N-terminal kinase (JNK) signaling pathway. 1-3 respectively indicate transcripts significantly ($p < 0.05$) altered by the low nutrient (LN) diet, AOX expression and the LN-AOX interaction, according to a multifactorial analysis performed using RNA-seq data and the software edgeR[50]. The columns LN, AOX and LN-AOX show respectively color representations of the $log_2$(fold change) values of the indicated transcript in control larvae cultured on LN diet, AOX-expressing larvae cultured on standard (SD) diet, and AOX-expressing larvae culture on LN diet, each in a pairwise comparison with control larvae cultured on SD diet. Control larvae, progeny of *UAS-AOX$^{F6}$* and *w$^{1118}$*; AOX-expressing larvae, progeny of *UAS-AOX$^{F6}$* and *daGAL4*.

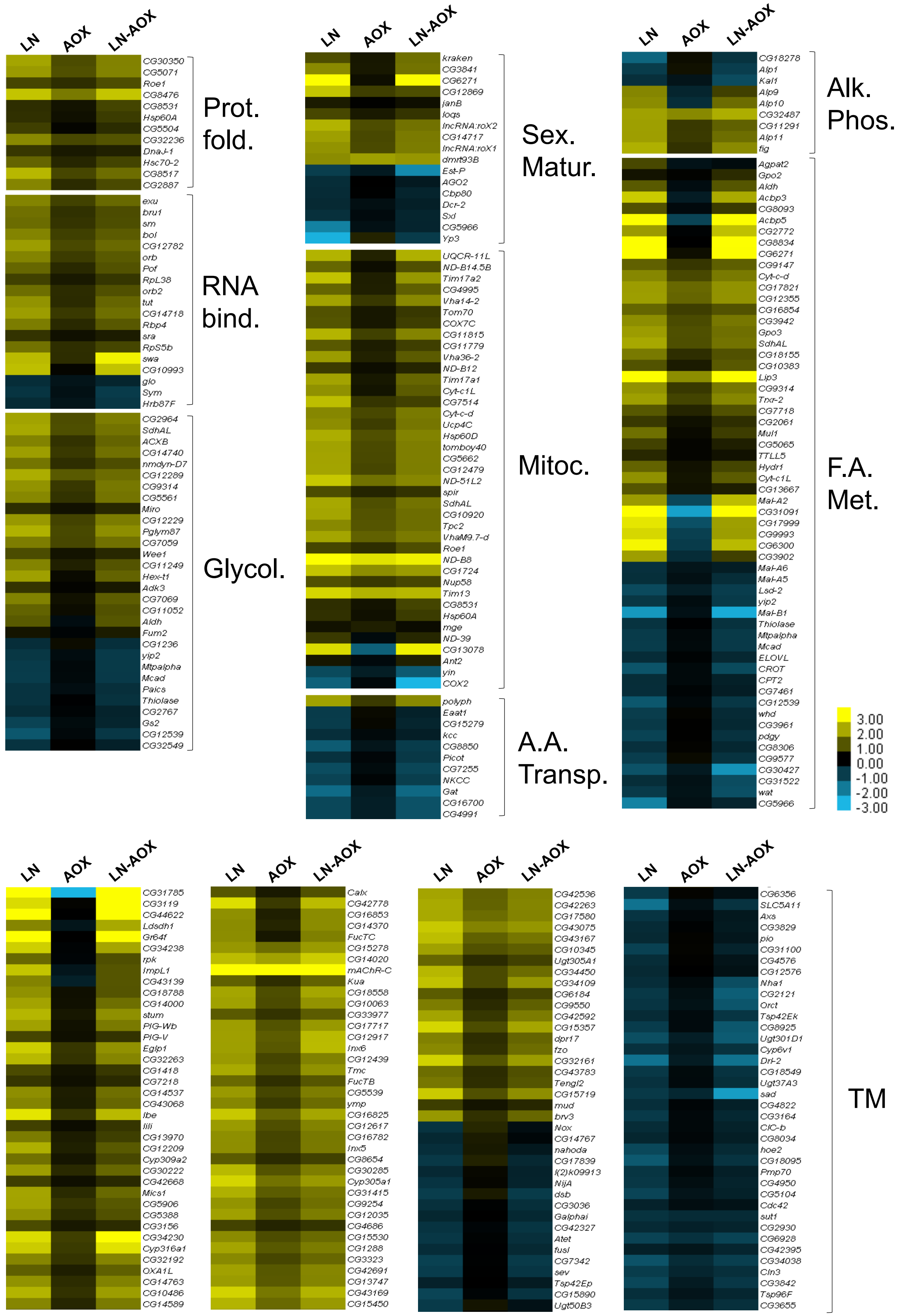
LN
AOX
LN-AOX
Prot. fold.
RNA bind.
Glycol.
Sex. Matur.
Mitoc.
A.A. Transp.
Alk. Phos.
F.A. Met.
TM
3.00
2.00
1.00
0.00
-1.00
-2.00
-3.00

**Figure S1.**

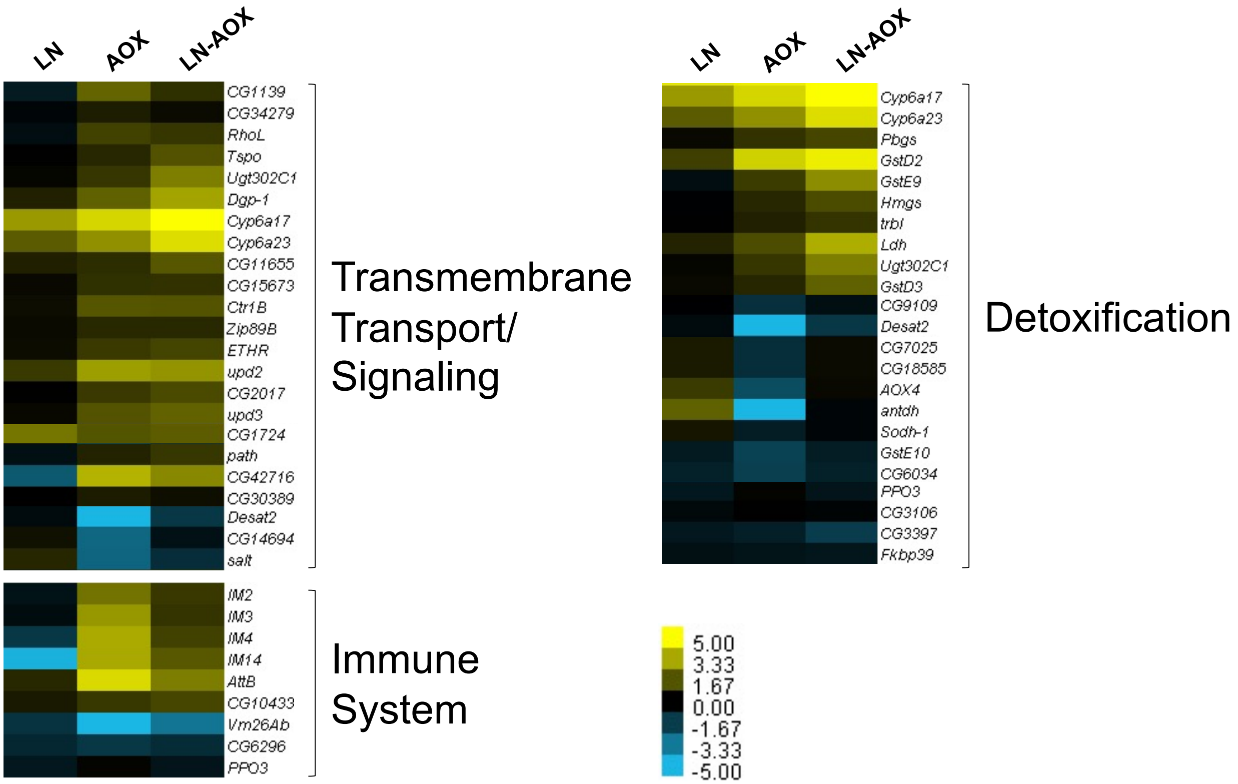
LN
AOX
LN-AOX
CG1139
CG34279
RhoL
Tspo
Ugt302C1
Dgp-1
Cyp6a17
Cyp6a23
CG11655
CG15673
Ctr1B
Zip89B
ETHR
upd2
CG2017
upd3
CG1724
path
CG42716
CG30389
Desat2
CG14694
salt
Transmembrane Transport/ Signaling
IM2
IM3
IM4
IM14
AttB
CG10433
Vm26Ab
CG6296
PPO3
Immune System
Cyp6a17
Cyp6a23
Pbgs
GstD2
GstE9
Hmgs
trbl
Ldh
Ugt302C1
GstD3
CG9109
Desat2
CG7025
CG18585
AOX4
antdh
Sodh-1
GstE10
CG6034
PPO3
CG3106
CG3397
Fkbp39
Detoxification
5.00
3.33
1.67
0.00
-1.67
-3.33
-5.00

**Figure S2.**

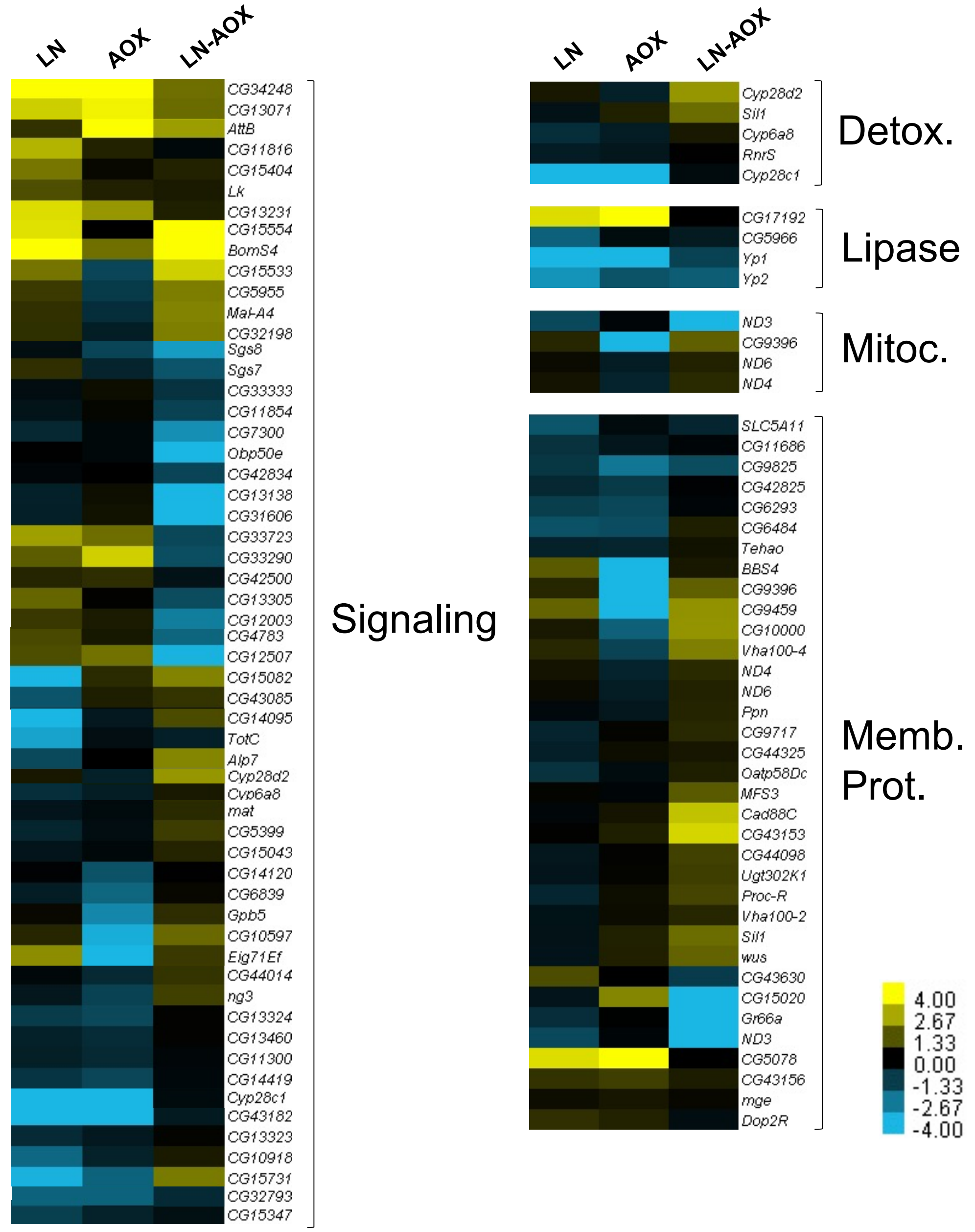
LN
AOX
LN-AOX
CG34248
CG13071
AttB
CG11816
CG15404
Lk
CG13231
CG15554
BomS4
CG15533
CG5955
Mal-A4
CG32198
Sgs8
Sgs7
CG33333
CG11854
CG7300
Obp50e
CG42834
CG13138
CG31606
CG33723
CG33290
CG42500
CG13305
CG12003
CG4783
CG12507
CG15082
CG43085
CG14095
TotC
Alp7
Cyp28d2
Cyp6a8
mat
CG5399
CG15043
CG14120
CG6839
Gpb5
CG10597
Eig71Ef
CG44014
ng3
CG13324
CG13460
CG11300
CG14419
Cyp28c1
CG43182
CG13323
CG10918
CG15731
CG32793
CG15347
Signaling
LN
AOX
LN-AOX
Cyp28d2
Sil1
Cyp6a8
RnrS
Cyp28c1
Detox.
CG17192
CG5966
Yp1
Yp2
Lipase
ND3
CG9396
ND6
ND4
Mitoc.
SLC5A11
CG11686
CG9825
CG42825
CG6293
CG6484
Tehao
BBS4
CG9396
CG9459
CG10000
Vha100-4
ND4
ND6
Ppn
CG9717
CG44325
Oatp58Dc
MFS3
Cad88C
CG43153
CG44098
Ugt302K1
Proc-R
Vha100-2
Sil1
wus
CG43630
CG15020
Gr66a
ND3
CG5078
CG43156
mge
Dop2R
Memb.
Prot.
4.00
2.67
1.33
0.00
-1.33
-2.67
-4.00

**Figure S3.**

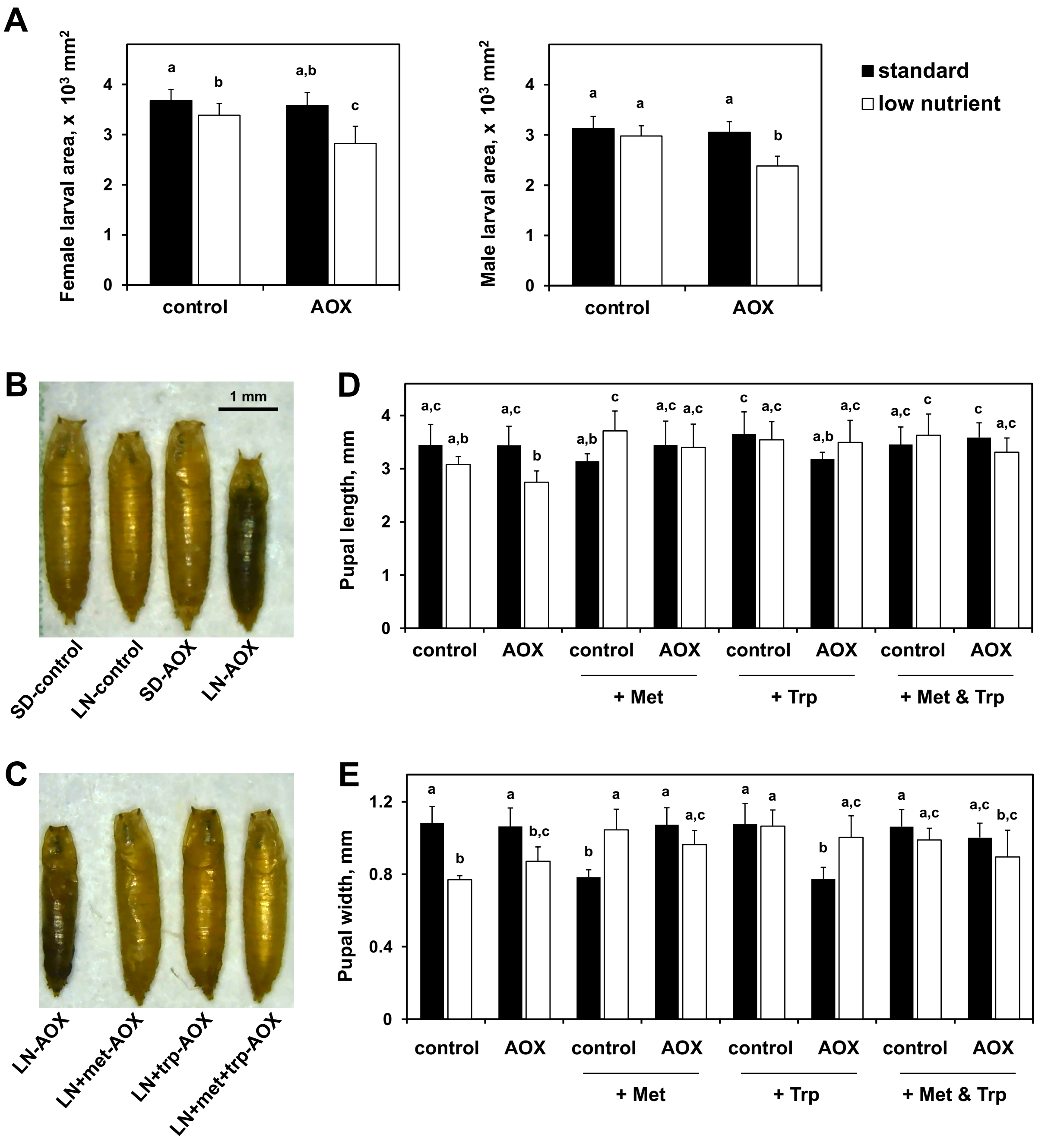
A
Female larval area, x 10^3 mm^2
Male larval area, x 10^3 mm^2
standard
low nutrient
control
AOX
B
1 mm
SD-control
LN-control
SD-AOX
LN-AOX
C
LN-AOX
LN+met-AOX
LN+trp-AOX
LN+met+trp-AOX
D
Pupal length, mm
+ Met
+ Trp
+ Met & Trp
E
Pupal width, mm

Figure S4.

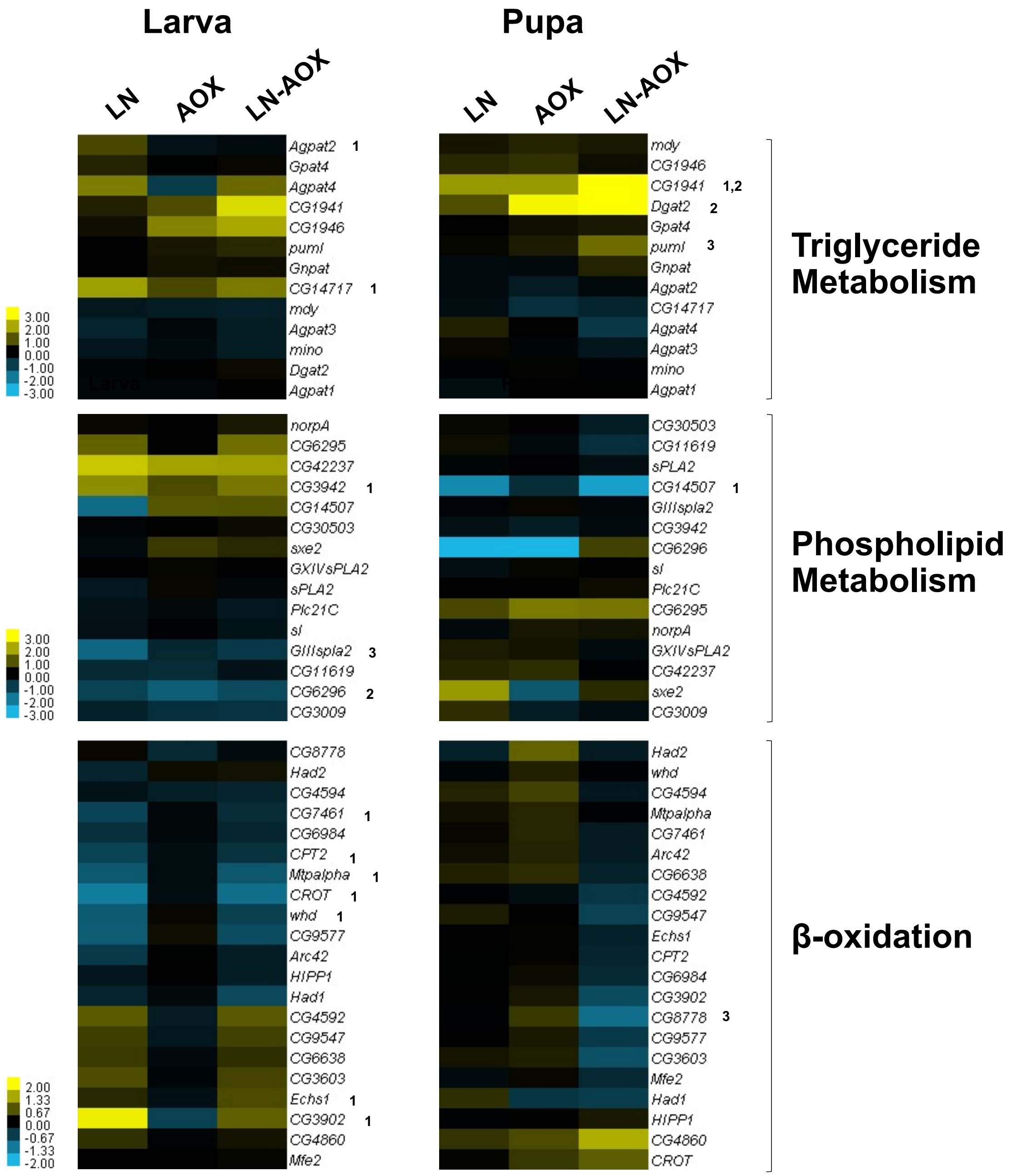
Larva
Pupa
LN
AOX
LN-AOX
Triglyceride Metabolism
Phospholipid Metabolism
β-oxidation
Agpat2 1
Gpat4
Agpat4
CG1941
CG1946
puml
Gnpat
CG14717 1
mdy
Agpat3
mino
Dgat2
Agpat1
mdy
CG1946
CG1941 1,2
Dgat2 2
Gpat4
puml 3
Gnpat
Agpat2
CG14717
Agpat4
Agpat3
mino
Agpat1
3.00
2.00
1.00
0.00
-1.00
-2.00
-3.00
norpA
CG6295
CG42237
CG3942 1
CG14507
CG30503
sxe2
GXIVsPLA2
sPLA2
Plc21C
sl
GIIIspla2 3
CG11619
CG6296 2
CG3009
CG30503
CG11619
sPLA2
CG14507 1
GIIIspla2
CG3942
CG6296
sl
Plc21C
CG6295
norpA
GXIVsPLA2
CG42237
sxe2
CG3009
CG8778
Had2
CG4594
CG7461 1
CG6984
CPT2 1
Mtpalpha 1
CROT 1
whd 1
CG9577
Arc42
HIPP1
Had1
CG4592
CG9547
CG6638
CG3603
Echs1 1
CG3902 1
CG4860
Mfe2
Had2
whd
CG4594
Mtpalpha
CG7461
Arc42
CG6638
CG4592
CG9547
Echs1
CPT2
CG6984
CG3902
CG8778 3
CG9577
CG3603
Mfe2
Had1
HIPP1
CG4860
CROT
2.00
1.33
0.67
0.00
-0.67
-1.33
-2.00

**Figure S5.**

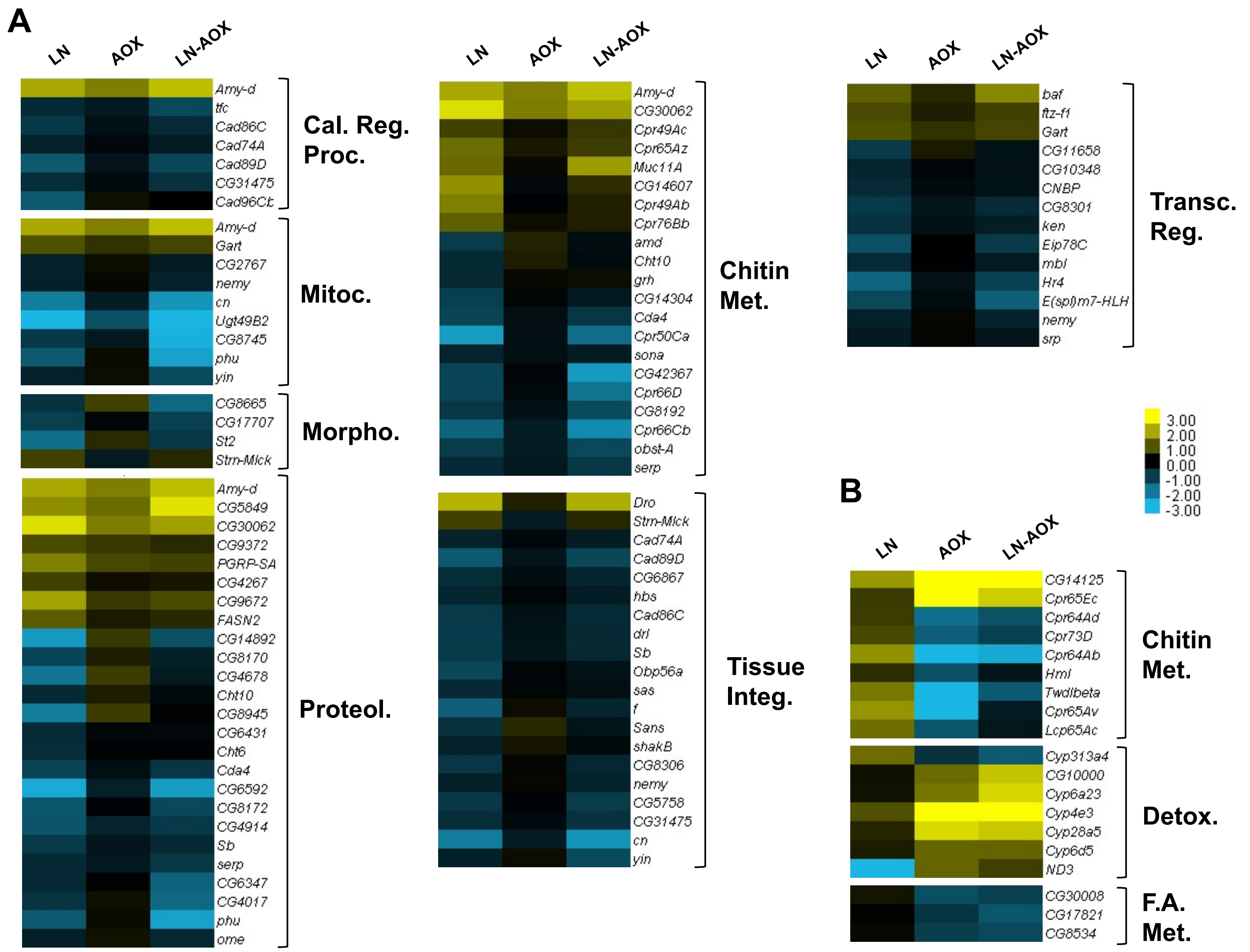
A
LN
AOX
LN-AOX
Cal. Reg. Proc.
Mitoc.
Morpho.
Proteol.
Chitin Met.
Tissue Integ.
Transc. Reg.
B
Chitin Met.
Detox.
F.A. Met.
3.00
2.00
1.00
0.00
-1.00
-2.00
-3.00

**Figure S6.**

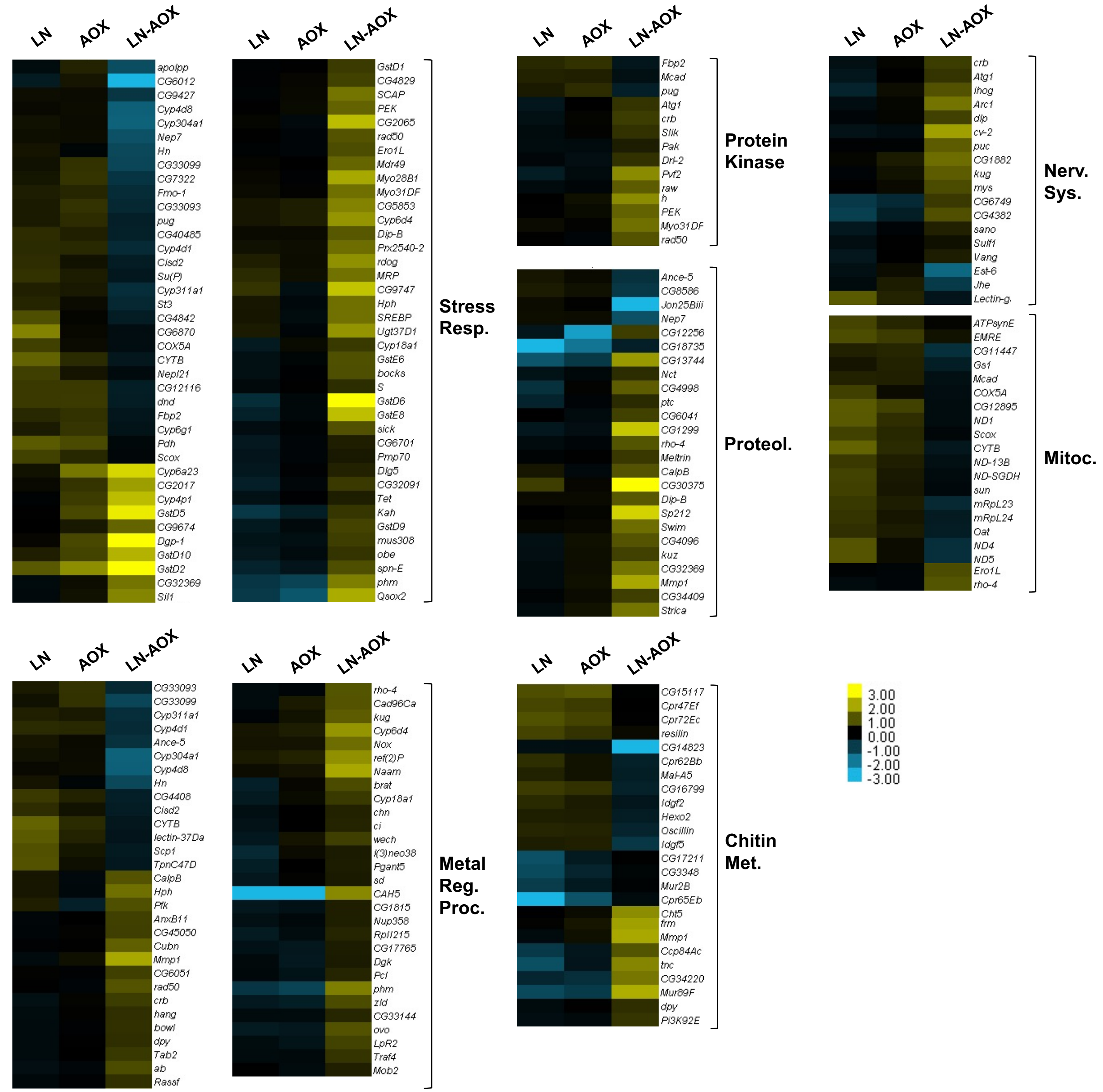
LN
AOX
LN-AOX
apolpp
CG6012
CG9427
Cyp4d8
Cyp304a1
Nep7
Hn
CG33099
CG7322
Fmo-1
CG33093
pug
CG40485
Cyp4d1
Cisd2
Su(P)
Cyp311a1
St3
CG4842
CG6870
COX5A
CYTB
Nepl21
CG12116
dnd
Fbp2
Cyp6g1
Pdh
Scox
Cyp6a23
CG2017
Cyp4p1
GstD5
CG9674
Dgp-1
GstD10
GstD2
CG32369
Sil1
GstD1
CG4829
SCAP
PEK
CG2065
rad50
Ero1L
Mdr49
Myo28B1
Myo31DF
CG5853
Cyp6d4
Dip-B
Prx2540-2
rdog
MRP
CG9747
Hph
SREBP
Ugt37D1
Cyp18a1
GstE6
bocks
S
GstD6
GstE8
sick
CG6701
Pmp70
Dlg5
CG32091
Tet
Kah
GstD9
mus308
obe
spn-E
phm
Qsox2
Stress Resp.
Fbp2
Mcad
pug
Atg1
crb
Slik
Pak
Drl-2
Pvf2
raw
h
PEK
Myo31DF
rad50
Protein Kinase
Ance-5
CG8586
Jon25Biii
Nep7
CG12256
CG18735
CG13744
Nct
CG4998
ptc
CG6041
CG1299
rho-4
Meltrin
CalpB
CG30375
Dip-B
Sp212
Swim
CG4096
kuz
CG32369
Mmp1
CG34409
Strica
Proteol.
crb
Atg1
ihog
Arc1
dlp
cv-2
puc
CG1882
kug
mys
CG6749
CG4382
sano
Sulf1
Vang
Est-6
Jhe
Lectin-g.
Nerv. Sys.
ATPsynE
EMRE
CG11447
Gs1
Mcad
COX5A
CG12895
ND1
Scox
CYTB
ND-13B
ND-SGDH
sun
mRpL23
mRpL24
Oat
ND4
ND5
Ero1L
rho-4
Mitoc.
CG33093
CG33099
Cyp311a1
Cyp4d1
Ance-5
Cyp304a1
Cyp4d8
Hn
CG4408
Cisd2
CYTB
lectin-37Da
Scp1
TpnC47D
CalpB
Hph
Pfk
AnxB11
CG45050
Cubn
Mmp1
CG6051
rad50
crb
hang
bowl
dpy
Tab2
ab
Rassf
rho-4
Cad96Ca
kug
Cyp6d4
Nox
ref(2)P
Naam
brat
Cyp18a1
chn
ci
wech
l(3)neo38
Pgant5
sd
CAH5
CG1815
Nup358
Rpl1215
CG17765
Dgk
Pcl
phm
zld
CG33144
ovo
LpR2
Traf4
Mob2
Metal Reg. Proc.
CG15117
Cpr47Ef
Cpr72Ec
resilin
CG14823
Cpr62Bb
Mal-A5
CG16799
Idgf2
Hexo2
Oscillin
Idgf5
CG17211
CG3348
Mur2B
Cpr65Eb
Cht5
frm
Mmp1
Ccp84Ac
tnc
CG34220
Mur89F
dpy
Pi3K92E
Chitin Met.
3.00
2.00
1.00
0.00
-1.00
-2.00
-3.00

**Figure S7.**

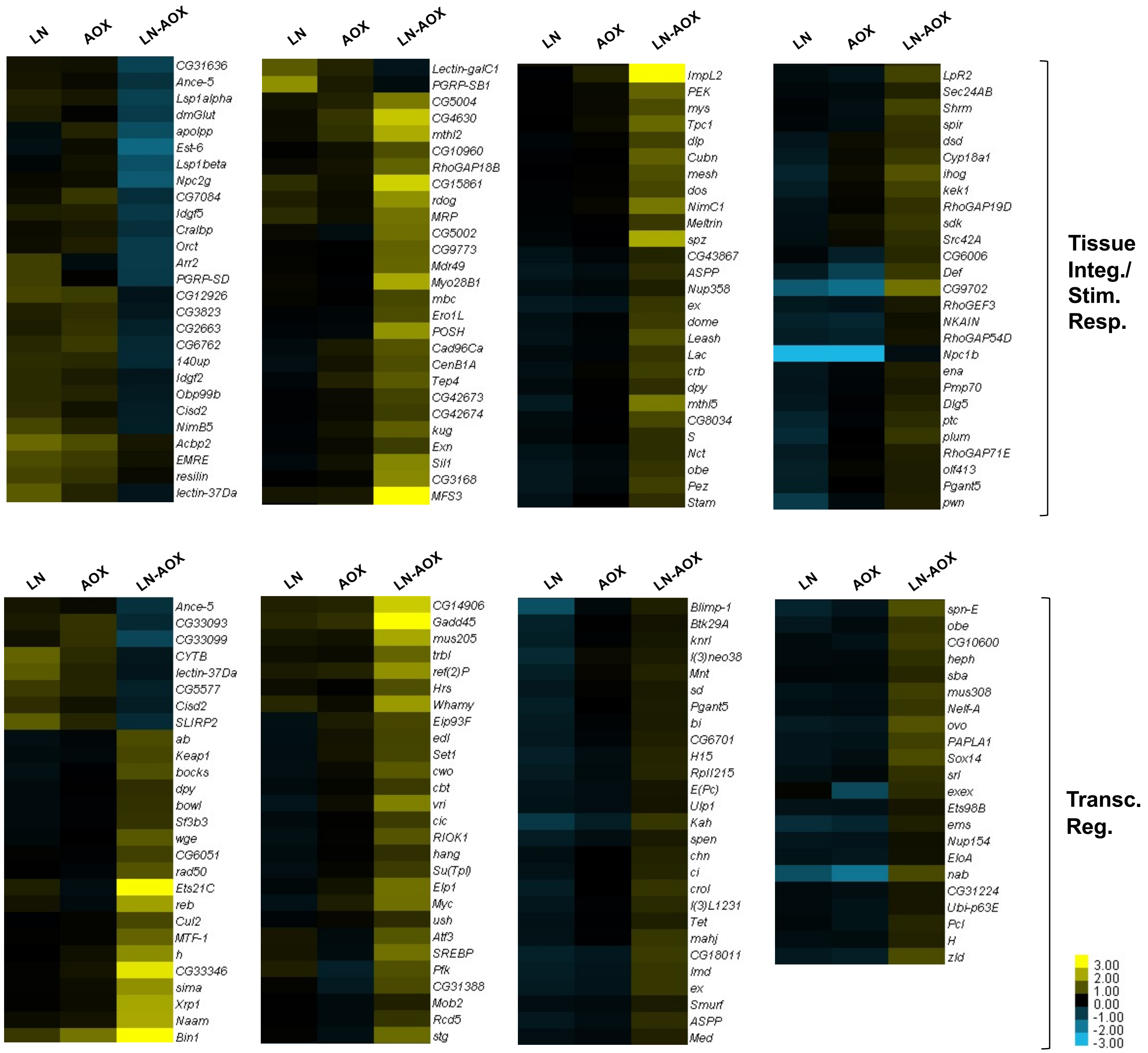


**Figure S7 (continued).**

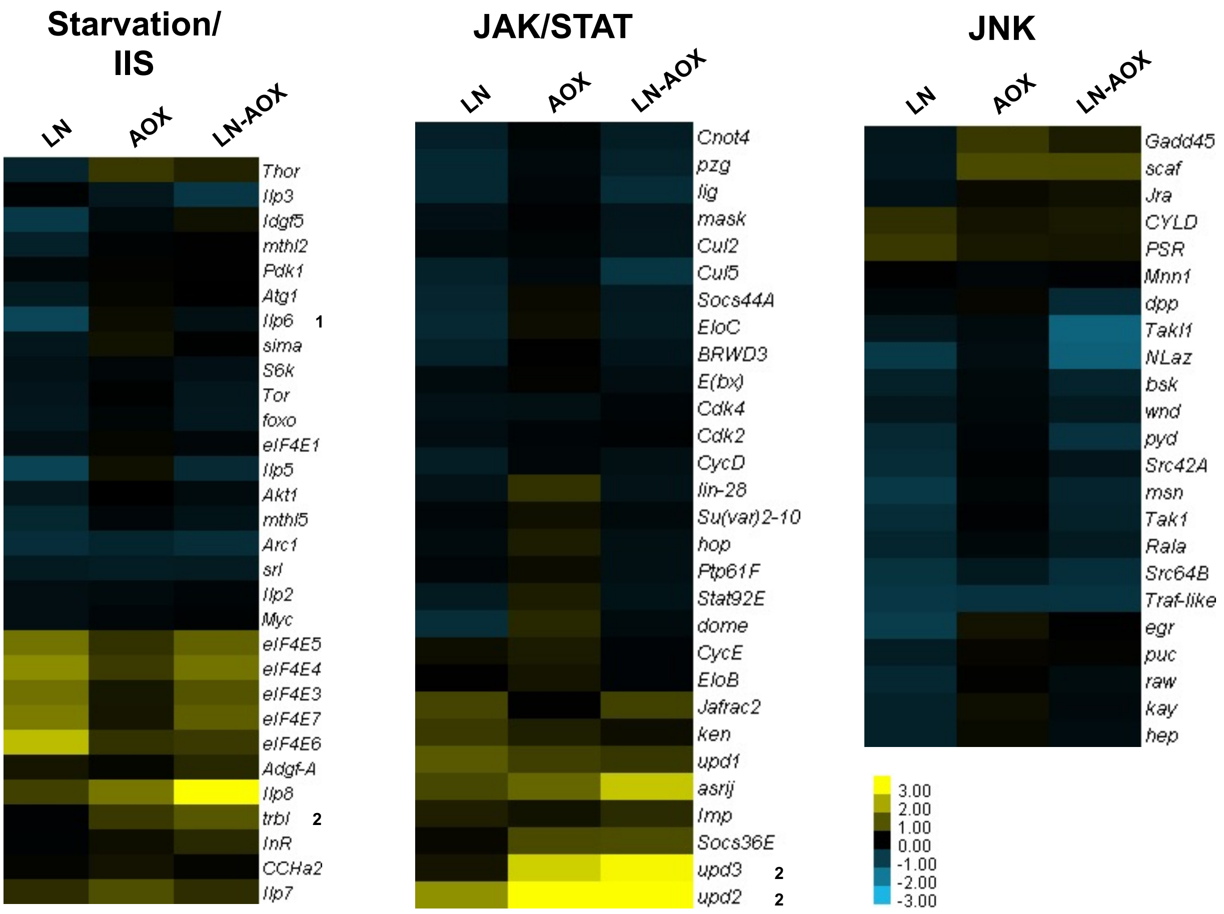

Starvation/
IIS
LN
AOX
LN-AOX
Thor
Ilp3
Idgf5
mthl2
Pdk1
Atg1
Ilp6 1
sima
S6k
Tor
foxo
eIF4E1
Ilp5
Akt1
mthl5
Arc1
srl
Ilp2
Myc
eIF4E5
eIF4E4
eIF4E3
eIF4E7
eIF4E6
Adgf-A
Ilp8
trbl 2
InR
CCHa2
Ilp7
JAK/STAT
LN
AOX
LN-AOX
Cnot4
pzg
lig
mask
Cul2
Cul5
Socs44A
EloC
BRWD3
E(bx)
Cdk4
Cdk2
CycD
lin-28
Su(var)2-10
hop
Ptp61F
Stat92E
dome
CycE
EloB
Jafrac2
ken
upd1
asrij
Imp
Socs36E
upd3 2
upd2 2
JNK
LN
AOX
LN-AOX
Gadd45
scaf
Jra
CYLD
PSR
Mnn1
dpp
Takl1
NLaz
bsk
wnd
pyd
Src42A
msn
Tak1
Rala
Src64B
Traf-like
egr
puc
raw
kay
hep
3.00
2.00
1.00
0.00
-1.00
-2.00
-3.00


**Figure S8.**
